# At the Southern Limit: The Arenal Central Site and the Guaraní Occupation of the Paraná Delta (Argentina)

**DOI:** 10.1101/2025.04.21.648367

**Authors:** Daniel Loponte, Mirian Carbonera, Fernanda Schneider, Neli Machado, Aline Bertoncello, Maicon Telles Szczygel, Alejandro Acosta, Barbara Mazza, Sheila Alí, Isabel Capparelli

## Abstract

This study analyzes the archaeological record of the Arenal Central site, a Guaraní residential base located on Martín García Island, within the Río de la Plata estuary—marking the southernmost extent of this population originating from the tropical forests of South America. The results significantly advance our understanding of these Amazonian forager-horticulturalists in the region, allowing us to explore a range of strategies and material culture expressions developed in an environment suboptimal for canoe-based populations. The study also presents a new set of chronological data, substantially increasing the number of radiocarbon dates available for these groups in the area. Findings indicate that Arenal Central was occupied around 1400 CE, during the final phase of the Guaraní expansion across the La Plata Basin. From this location, a vast catchment area was incorporated, including the lower Uruguay River, the Paraná Delta islands, and the adjacent Uruguayan steppe. The extensive use of territory appears to have been a strategic response to the island’s limited carrying capacity and the scarcity of key resources required to sustain the Guaraní ecological niche. While the recovered assemblage preserves the core features of Guaraní material culture, a marked decline is observed in the complexity of pictorial expressions—particularly in painted ceramics. This simplification may reflect reduced intra-ethnic interaction, likely due to the low density of Guaraní populations in the surrounding area, as well as the challenges imposed by a constrained insular setting. We also compare subsistence strategies and ceramic assemblages from other sites that have been excavated and analyzed following comparable standards. Finally, the results are contextualized within the broader process of Guaraní occupation of the Paraná Delta and the Río de la Plata estuary.

## 1. Introduction

This study aims to report the principal results of the 2023 excavation season at the Arenal Central site, located on Martín García Island in the upper estuary of the Río de la Plata, southeastern South America. This site was occupied by Amazonian forager-horticulturalists whose material culture is classified within the Guaraní Archaeological Unit (Section 2). Originating in the southern Amazon basin, this population expanded across southeastern South America, covering over 1,500 km from southern Brazil to the Río de la Plata estuary in Argentina, likely following the Uruguay River as their primary migratory route to this region (Figure 1 and Section 2) (Brochado, 1984, 1989; Schmitz, 1991; Noelli, 1999-2000; Bonomo et al., 2015; Noelia & Correa, 2016; López Mazz & López Cabral, 2020; Loponte et al., 2025, among others).

**Figure 1.**
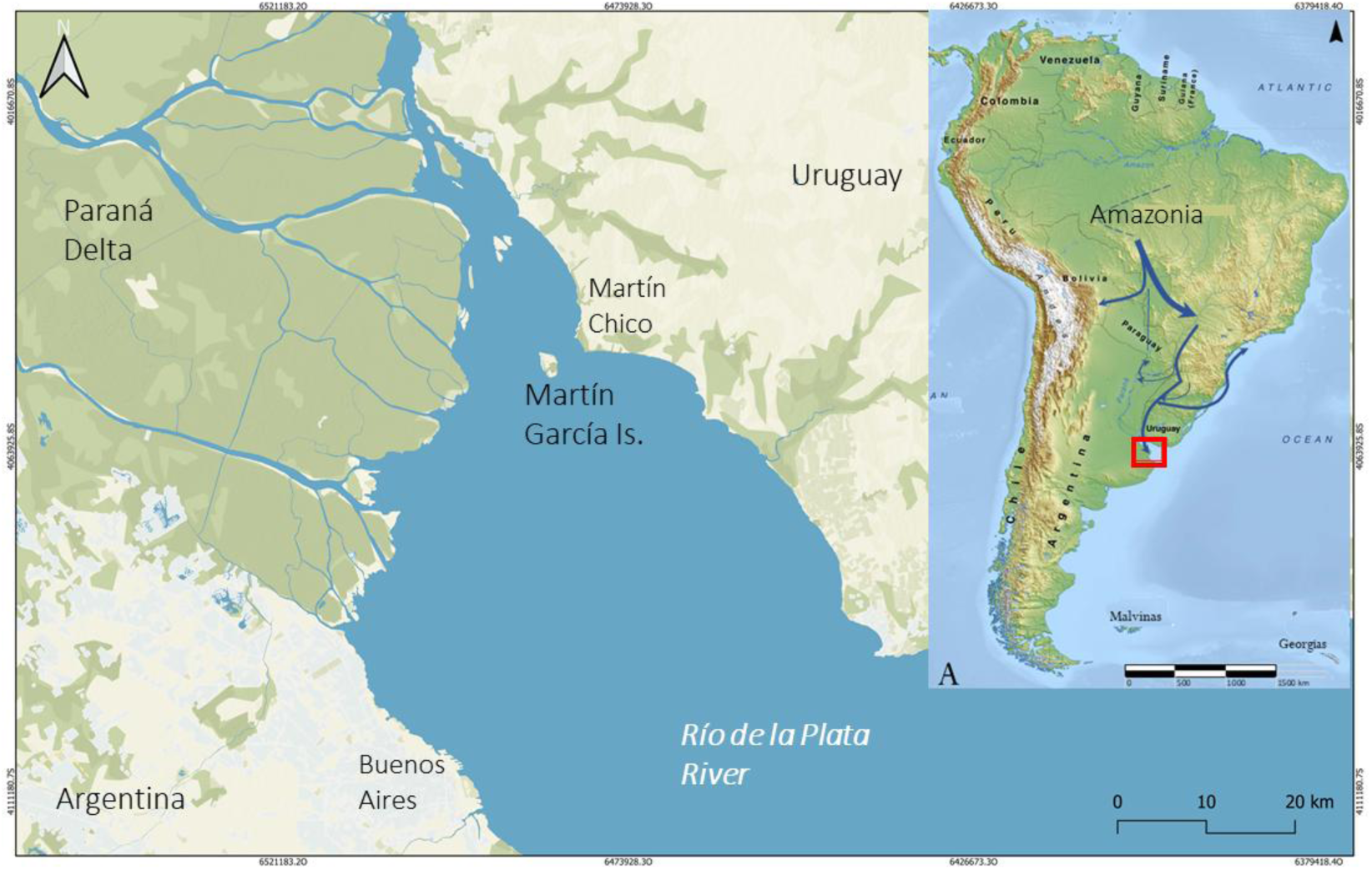
Location of Martín García Island within the Estuary of the Río de la Plata. In the right and smaller map, the expansion of Guaraní population from Southwest Amazonia to the Río de la Plata. Martín García Island is enclosed in a red square

The archaeological record of Arenal Central is particularly noteworthy as it represents the southernmost settlement of this population along the Uruguay River axis. Its significance is further enhanced by the island’s pronounced insularity, small size, and its location in an environment vastly different from the population’s original homeland. The region surrounding Arenal Central integrates tropical forests with Chaco and Pampas-like ecosystems, creating a unique ecological setting that combines elements of these diverse biomes. As a result, the site’s archaeological record provides valuable insights into the cultural variability of these groups as they adapted to environments beyond the tropical and subtropical forests that constituted their traditional habitat.

To achieve the objectives of this study, we first describe the site, highlighting key aspects of its stratigraphy and formation processes, along with five new radiocarbon dates that refine the chronology of the Guaraní occupation on the island. We then analyze the composition of the faunal, lithic, and ceramic assemblages, examining the catchment areas for subsistence resources and raw materials. Additionally, we discuss geomorphological changes in the area driven by the progradation of the Paraná Delta, which have altered resource availability since the site’s occupation. Finally, we compare these findings with the best-documented records from the region and other relevant datasets^1^, providing a broader comparison of key analytical categories in this study and an overview of Guaraní colonization in the area.

## 2. Archaeological background

The Guaraní archaeological record reflects the material culture and behavioral patterns associated with the lifeways of Amazonian forager-horticulturalist populations who migrated from southwestern Amazonia to the Plata Basin during the Late Holocene, in pre-Columbian times (Brochado 1973, 1984, 1989; Noelli 1999–2000; Schmitz 1991, among others).Their subsistence relied on hunting, fishing, gathering, and cultivating crops such as maize, tropical tubers, beans, squash, and peanuts, among others. These groups also practiced ritual anthropophagy (Ramírez [1528] in Madero, 1939; Fernández de Oviedo y Valdés [ca. 1526–1557] 1945; Schmidl [ca. 1536–1554] 1948). Although the chronology, geographic extent, and cultural processes associated with the emergence of these assemblages in southwestern Amazonia remain uncertain, it is worth noting that some stylistic features characteristic of their pottery and other artifacts appear to have been present in that region as early as 2000 BCE (Almeida, 2013; Caldarelli, 2008; Miller, 2009; Zimpel, 2009, 2018, among others). Therefore, it is correct to consider that the Guaraní expansion—explicitly referred to as such—originated in the southwestern Amazon, as illustrated by the smaller map in Figure 1 (see also Brochado 1973, 1984; Zimpel, 2009, 2018). Guaraní archaeological assemblages are well-defined by consistent associations of material culture traits and specific practices. Pottery is especially distinctive, including corrugated and polychrome vessels with emblematic geometric motifs—mainly in red, orange, brown, and black—often applied over white slips. Vessel typologies are highly standardized, correlating with functional and emic categories. The assemblage also features characteristic ornaments such as T-shaped quartz labrets, gourd-shaped pendants, bone and stone earrings, and clay pipes. Common tools include square, neckless axes and whetstones, likely used for maintenance and crafting. Mortuary practices involved both primary and secondary burials, often in urns, with or without grave goods, reflecting broader Amazonian traditions. The lithic assemblages are dominated by small flakes of microcrystalline rocks. Quadrangular axe heads without necks are also frequently found (Ambrosetti, 1895, Ali et al., 2017; Brochado 1973, 1984; Brochado & Monticelli, 1994; Buc, 2017; Capparelli, 2017; Carbonera, 2014; Carbonera & Loponte, 2024, 2015; Carbonera et al., 2014, 2021; La Salvia & Brochado, 1989; Lothrop, 1932; Maldonado Bruzzone, 1931; Mazza et al., 2016; Milheira 2014; Muller & Souza, 2011; Musali, 2010a; Novasco et al., 2021; Noelli, 1993, 1999; Pérez & Alí, 2017; Prous, 2010; Prous & Lima, 2008; Rogge, 1996; Schmitz, 1991, 2008; Schneider et al., 2014a, 2014b; Sempé & Caggiano, 1995; Silvestre & Capparelli, 2017, among many others).

In the La Plata Basin, the earliest Guaraní sites date around 500 ± 100 CE, corresponding to settlements located in the Upper Paraná River. From these areas, the Guaraní population expanded eastward, reaching the Atlantic coast of southern Brazil around 800-1000 CE, and the Paraná Delta and Río de la Plata estuary between 1229 and 1423 CE (Loponte & Acosta, 2003-2005) coinciding with their maximum territorial extent (Figure 1). This expansion appears to have been driven by the colonization of new territories by founder populations with low demographic densities, establishing new settlement areas while leaving intermediate zones uncolonized, likely inhabited by pre-existing populations. These new population centers gradually expanded by saturating the adjacent space and creating additional colonization areas.

The Guaraní expansion along the south axis of South America was facilitated in part by canoe mobility along the rivers within a tropical forest environment, which provided ideal conditions for cultivating most of their tropical-origin crops and supported faunal resources these groups exploited. These forests also contained a rich diversity of plant species used for food, cosmetics, medicine, rituals, and other raw materials, all of which contributed to the development of the Guaraní ecological niche (e.g., Acosta et al., 2019; Brochado, 1984; Noelli, 1993, 1999; Schmitz, 1991; Schneider et al., 2024a, among others).

Archaeological research on Guaraní sites in the Río de la Plata estuary and Paraná Delta has a long, though limited, academic tradition. These studies began with the work of Outes (1917), based on a few materials recovered by Antonio Pozzi from the Puerto Viejo site on Martín García Island, and from Arroyo Largo in the Paraná Delta, unsystematically collected by Enrique de Carles (Outes, 1918). In the first half of the 20th century, additional surface collections and limited interventions were carried out by local museum curators and amateur collectors (e.g., Vignati, 1936; Bonomo et al., 2009; Pazzi, 2021; Torino & Bonomo, 2024). During this period, curators from the Museo de La Plata recovered human remains from the Arroyo Fredes and Arroyo Malo sites (Vignati, 1941), which were recently analyzed with a focus on mortuary practices and diet (Loponte et al., 2016; Mazza et al., 2016).

Excavations at Guaraní sites conducted in accordance with modern academic standards began in the 1930s with Lothrop’s (1932) investigations at Arroyo Malo. These initial studies offered valuable insights into the burial patterns observed at the site, along with examples of material culture recovered from the excavated area, which appears to have been used almost exclusively for funerary purposes.

Subsequent efforts included limited fieldwork conducted by Cigliano (1968) at the El Arbolito site on Martín García Island and by Caggiano (1982) at the Paraná Guazú 3 site in the Paraná Delta. Both researchers published only limited results from their investigations, including a radiocarbon age for the El Arbolito site, marking the first such date obtained for a Guaraní site in the area, which remained the sole radiocarbon determination until 2003–2005 (see below). In a manuscript report on the Paraná Guazú 3 site, Caggiano indicates that 4.5 m² were excavated, yielding 4,370 ceramic fragments but no faunal remains. This absence is particularly striking given the excellent conditions for bone preservation in the area. The large amount of painted pottery is also particularly noteworthy (see Section 6.4) which was analyzed by and Pérez (2016) and Pérez & Alí (2017).

In the 21^st^ century, research activities resumed with systematic excavations by Loponte and Acosta (2003–2005) at the Arroyo Fredes site (Paraná Delta) and by Capparelli (2014) at Arenal Central on Martín García Island. These recent projects represent systematic investigations of well-preserved Guaraní residential areas, generating a substantial body of new data and providing fresh perspectives on previously unexplored aspects of Guaraní archaeology in the region regarding material culture, funerary patterns, chronology, biological markers of activity, subsistence patterns and mobility (Acosta & Mucciolo, 2009; Acosta et al., 2010, 2019; Alí et al., 2017; Capparelli, 2014, 2019; Buc, 2017; Buc et al., 2014; Loponte et al., 2011, 2016; Mazza et al., 2016; Musali, 2010a; Pérez & Alí, 2017; Pérez et al., 2009, 2018; Silvestre & Buc, 2015; Silvestre & Capparelli, 2017).

In addition to the previously described efforts based on records adequately recovered, recent years have seen the unsystematic collection of Guaraní materials from heavily impacted sites such as Kirpach and Rincón de Milberg (Loponte & Capparelli, 2013, and unpublished data), as well as the discovery of what appear to be, for now, isolated burials on an island located along El Duraznito creek in the Paraná Delta (unpublished data). Starting in 2023, new research activities were initiated, introducing new topics for local Guaraní archaeology based on previously collected materials, along with the reevaluation of sites identified during the early stages of investigation, such as Arroyo Malo, Arroyo Largo, and Arroyo Fredes all located in the Paraná Delta (unpublished data). This new phase, driven by the incorporation of new researchers and funding resources, enables renewed efforts in excavation and analysis, marking a significant step forward in the development of Guaraní archaeology in the area, of which this work is a part.

## 3. Environmental settings

### 3.1. Geology and geomorphology

It is very important to describe the geomorphological characteristics of the island and its environmental conditions, as these aspects are scarcely documented in the archaeological literature and need to be introduced here to facilitate a better understanding of the text.

Martín García is a small rocky island with a surface area of approximately 2 km² and a maximum elevation of about 28 meters. It is located in the upper estuary of the Río de la Plata, near the mouth of the Uruguay River and separated from the mainland by the “Canal del Infierno,” a 3.5 km-wide channel, across which lies the Martín Chico area in the Eastern Republic of Uruguay (Figure 1). The island is an outcrop of the Precambrian crystalline basement, known as the Martín García igneous-metamorphic complex, mainly composed of granitoids, schists, orthogneisses, migmatites, ultrabasic rocks, and amphibolites. Among these, amphibolites—characterized by their granular texture and varying shades of green—are the most widespread rock type on the island (Dalla Salda, 1981; Benítez, 2023). In the central and northern sector, where the terrain forms a gentle plain, the bedrock is mostly covered by Quaternary sediments. However, in the southern area, with its more elevated and rugged topography, the bedrock emerges in narrow rocky strips that extend toward the southern and southeastern beaches, which are covered with blocks and clasts detached and eroded by fluvial activity (Figure 2).

**Figure 2.**
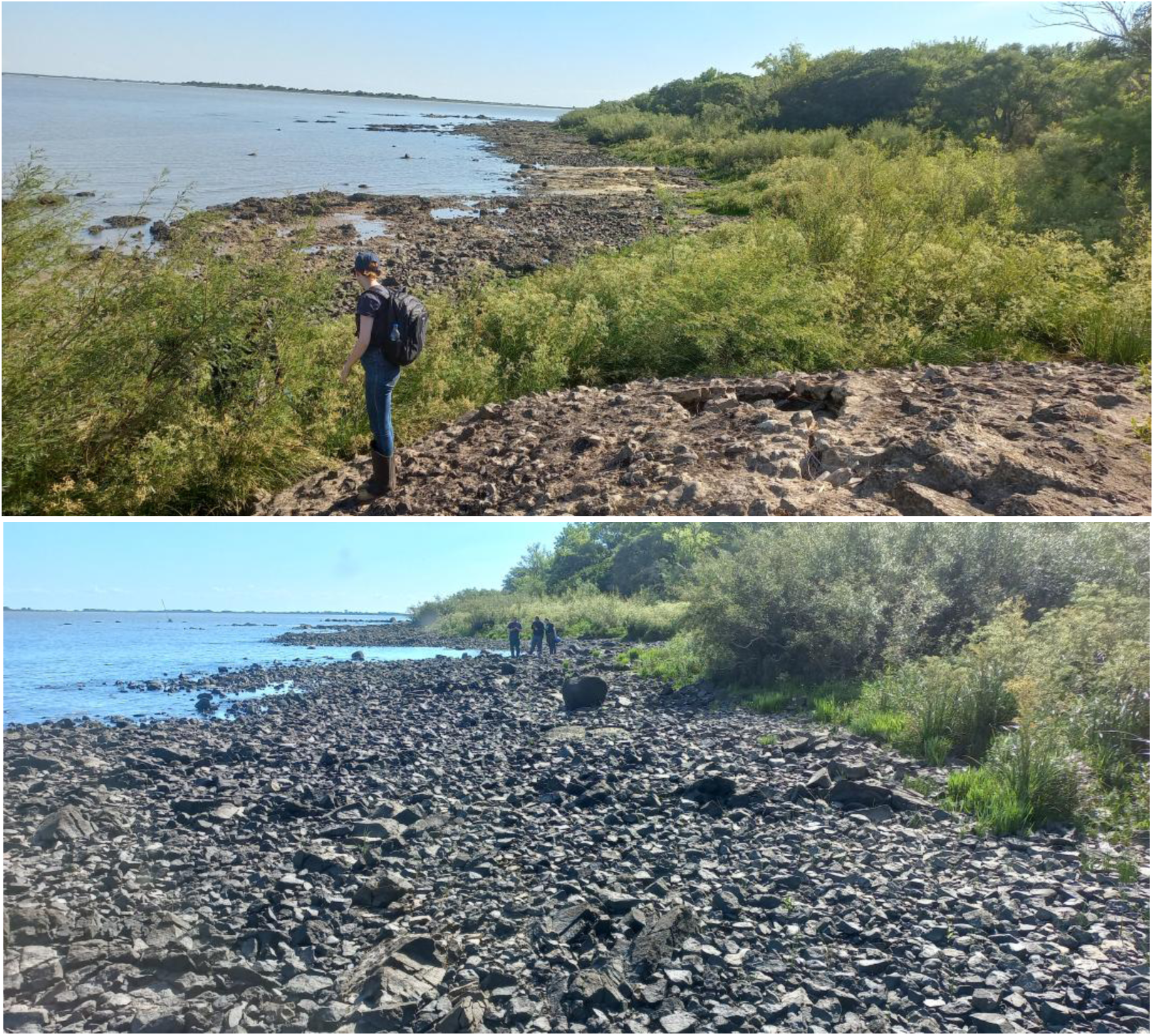
Southern beach of Martín García Island, covered with clasts and boulders. Photograph shows four of the authors of this study (legal disclaimer).

Across most of the island, the igneous-metamorphic complex is unconformably overlain by marine sediments known as “Unit A” (Ravizza, 1982), containing gravel ranging from sand to pebbles, predominantly composed of quartz, along with estuarine and marine gastropods and bivalves. Some profiles also display calcareous concretions. Unit A has been radiocarbon-dated to 19,050 ± 500 years ^14^C BP (Ravizza, 1982). Following this, continental aeolian silts of Pleistocene age (Unit B) were deposited, with thicknesses reaching up to 5 meters. Above these silts lies a 90 cm-thick sand layer corresponding to deposits from a mid-Holocene marine ingression (Unit C), dated to 5,740 ± 130 years ^14^C BP (Ravizza, 1982). Extensive accumulations of loose, unconsolidated sand, designated as Unit D, overlie Unit C. These deposits are particularly prominent in the central and western sectors of the island, forming the “Arenal Central” of Martín García. Although the origin of these sands remains uncertain, they may be linked to significant floods of the Uruguay River (Ravizza, 1982). Based on its stratigraphic position, Unit D is assigned to the Late Holocene (< 3,500 years BP).

The geomorphological configuration of the island has changed significantly over recent decades due to sediment transport and progradation, resulting in the formation of an extensive sedimentary area attached to its northern end, which began to emerge in the 1980s. This same process has filled the old bay located west of the Arenal Central, where Puerto Viejo area is situated, and the formation of new sedimentary island near-west of Martin García (Figure 3). This progradation process has also substantially reduced the distance between Martín García Island and the Paraná Delta, bringing them to just a few kilometers apart today. However, during the Guaraní occupation, the advancing front of the lower Delta was likely situated farther northwest, possibly 25 to 30 kilometers from its current position (Figure 3 and Section 7.3).

**Figure 3.**
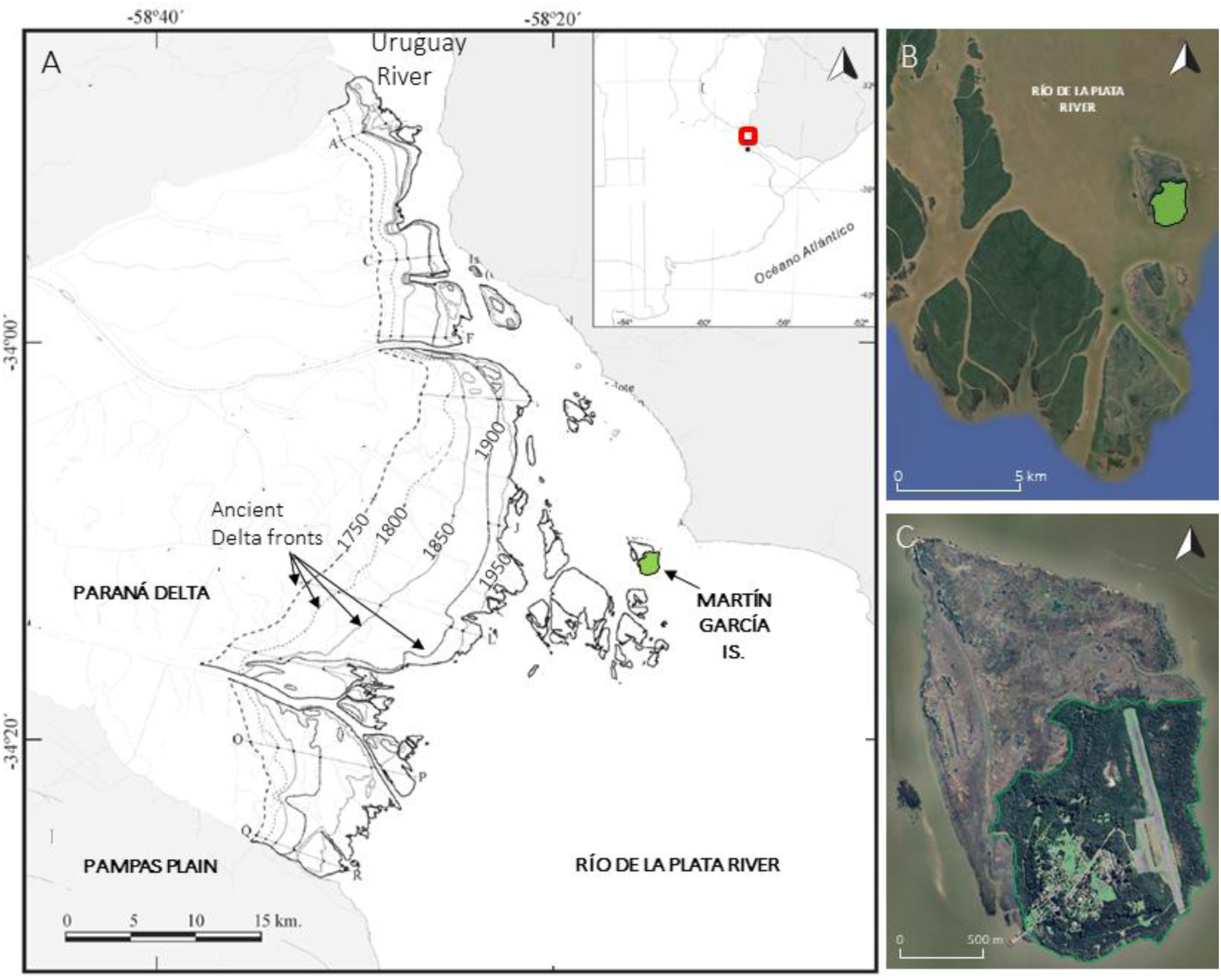
Map A: Location of Martín García Island in relation to recently formed sedimentary islands of the upper estuary of the Río de la Plata and the successive advancing fronts of the Paraná Delta, as documented in existing historical cartography (adapted and modified from Medina & Codignotto, 2013). B: Martín García Island (light green) alongside sedimentary islands to the north, west and south that emerged during the 20^th^ century. C: Detail of the original silhouette of Martín García Island, shown in light-dark green, with the sedimentary deposits attached to its northern and western sectors depicted in brown.

### 3.2. Environment

Martín García Island has a warm-temperate climate influenced by the subtropical regime of the Uruguay River, which extends these conditions into the upper estuary of the Río de la Plata. It is part of the Paraná wetland, a region dominated by wetlands and subtropical fluvial environments, including the islands of the Paraná Delta, adjacent continental floodplains, and the fluvial banks of the Uruguay and Paraná rivers. Ecologically, while the region is primarily dominated by tropical and subtropical species, it also exhibits significant influences from the adjacent Chaco-Pampean plains, which encompass temperate environments hosting both humid-temperate and xerophytic species. The latter vegetation grows particularly on coastal ridges left by the ingressive and regressive processes of the sea during the Holocene and on fluvial banks formed by loose sands that remained beyond the regular flood pulses (Burkart et al., 1999; Cabrera & Zardini, 1978).

Martín García Island exemplifies this complex interplay of ecosystems, reflecting the diversity and transitions between these distinct ecological zones. Along the beach-adjacent areas of the island, riparian forests thrive, featuring woody species such as *Salix humboldtiana* and *Erythrina crista-galli*, and several tree-like shrubs, such as *Sesbania punicea* and *Senna corymbose* adapted to subtropical environments and capable of withstanding brief periods of flooding. A short distance inland a denser forest emerges, representing a degraded extension of the Upper Paraná Atlantic Forest (also called “Paranaense Forest”; cf. Cabrera & Zardini, 1978). This multistratified forest canopy includes species such as *Rapanea* sp., *Ocotea acutifolia*, *Eugenia uruguayensis*, and *Syagrus romanzoffiana*. Beneath the canopy grow shrubs, bamboo species, and lianas, with a moss layer covering much of the forest floor. This formation is denser than the riparian forest, allowing significantly less light to reach the ground. Both forest types are found at elevations below 4 - 6 meters above sea level, forming a ring around the island’s terrain. It is worth noting that oceanic storms, which have a significant influence on the entire estuary, can raise water levels up to 3.6 meters, flooding areas below this altitude. Above this elevation, covering the island’s interior, a savanna environment and a xerophytic forest characteristic of the Tala District develops (Arturi and Juárez, 1997). This forest includes species such as *Scutia buxifolia*, *Celtis tala*, and *Jodina rhombifolia* (Arturi and Juárez, 1997), along with various cacti and shrubs that thrive in the sandy, porous soils of Units C and D. Areas where species from these three forest formations intermingle are common, and it is not unusual to find subtropical species growing within the xerophytic forest area. Based on the current distribution of plant communities, soil types, and the relationship between altitude and vegetation types on the island (e.g., Arturi and Juárez, 1997), it can be inferred that much of the island was covered by a tropical-subtropical forest assemblage, while its interior was occupied by a more open savanna interspersed with patches of xerophytic forest (Figure 4).

**Figure 4.**
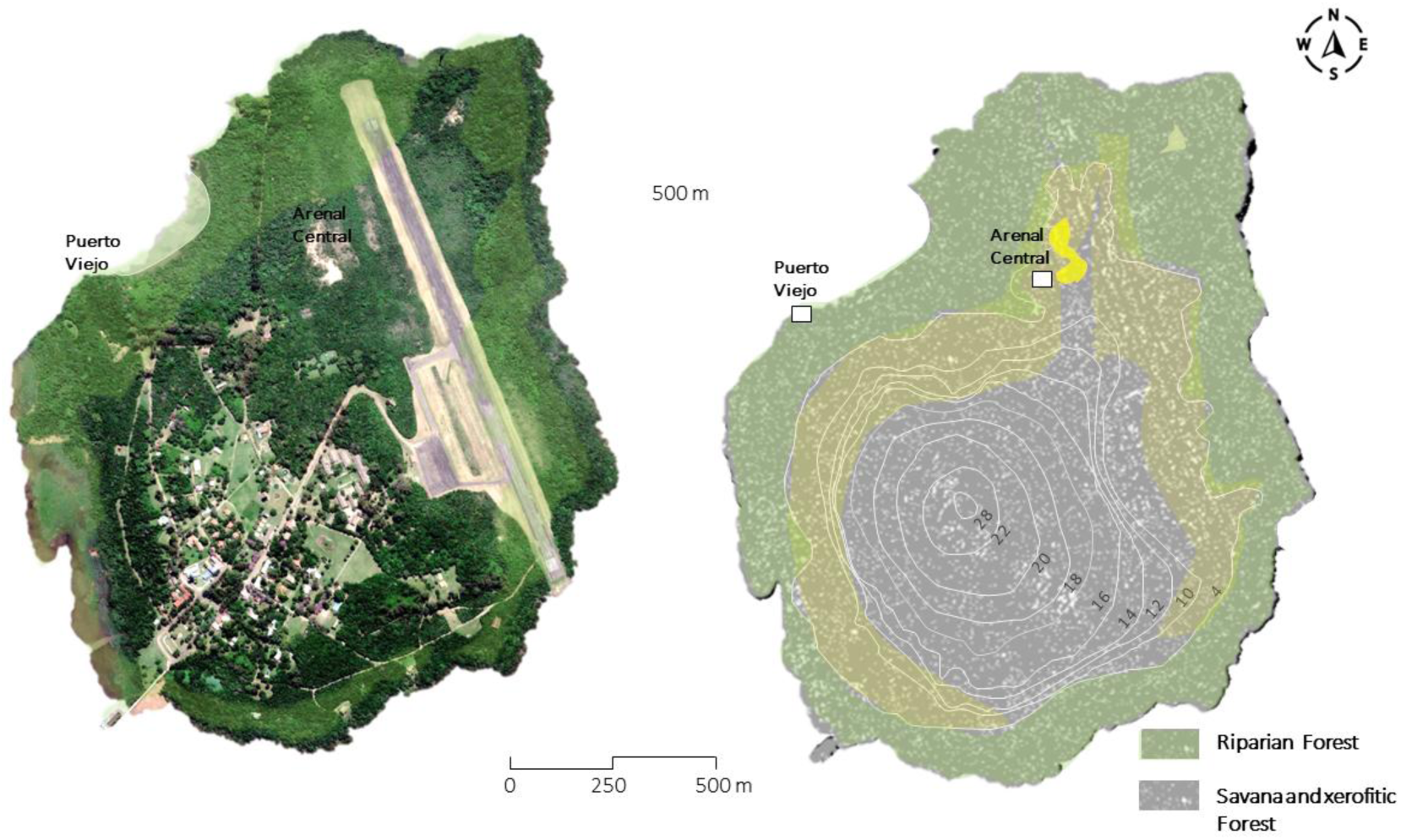
Left: Satellite image of Martín García Island displaying the locations of Guaraní archaeological sites, Puerto Viejo and Arenal Central. Right: hypothetical reconstruction of the areas occupied by the island’s plant communities during the Guaraní occupation, inferred from their present-day distribution and ground elevation. The overlapping-colored areas indicate sectors where marginal forest and xerophytic forest intersect. Note that the riparian forest includes the marginal forest. This image was specifically developed for this paper based on Dalla Salda (1981) elevation data (see other representations in Loponte et al., 2011, and Capparelli, 2019).

It is highly likely that, before the Guaraní occupation, the island hosted small populations of mammals typical of the area, such as *Blastocerus dichotomus* (marsh deer), *Myocastor coypus* (coypu), and *Hydrochoerus hydrochaeris* (capybara). The latter two species are excellent swimmers and adept colonizers of insular and aquatic environments. The marsh deer is also capable of swimming, although over shorter distances. In fact, during the 2023 excavations, an individual of this species reached the island by swimming. Species like this might also arrive at Martín García on floating islands composed of vegetation, which are common during major floods. These accumulations of floating vegetation are frequently carried along by the Paraná and Uruguay rivers and often include both small and large animals. However, the larger resident fauna was probably rapidly exploited and driven to extinction following the onset of Guaraní colonization of the island. This outcome can be attributed to the expected small population size of these local faunal communities and the pressures arising from resource exploitation by a Guaraní village, which typically consisted of dozens or even hundreds of people.

In contrast, the availability of fish resources around Martin García is remarkably abundant. The upper estuary serves as a natural extension of the fish populations from the Uruguay and Paraná rivers, where over 150 species have been documented, many of them large and exhibiting notable aggregation and migratory patterns (Loponte, 2008; Musali, 2010b). Just 3.5 km across the Del Infierno channel, on the Uruguayan mainland, lies a strip of riparian forest up to 1 km wide in the Martín Chico area. This forest then gives way to open grasslands, home to typical species such as *Rhea americana* (greater rhea), *Ozotoceros bezoarticus* (pampas deer), and dasypodids (armadillo species).

### 3.3. Location of the Arenal Central archaeological site

Arenal Central is situated on a gently sloping plain with xerophytic vegetation, spanning the central-northern sector of Martín García Island at elevations around 4 – 6 meters above sea level (Figure 4). The surface is covered by sand from Unit D, both loose and stabilized by vegetation, extending across the open dune field and the surrounding xerophytic forest (Figure 5). While the exact center of the site remains unknown, the excavation area featured in this study is located at latitude −34.180595° and longitude −58.250637° (± 2 m) (Figure 6; see also Section 5).

**Figure 5.**
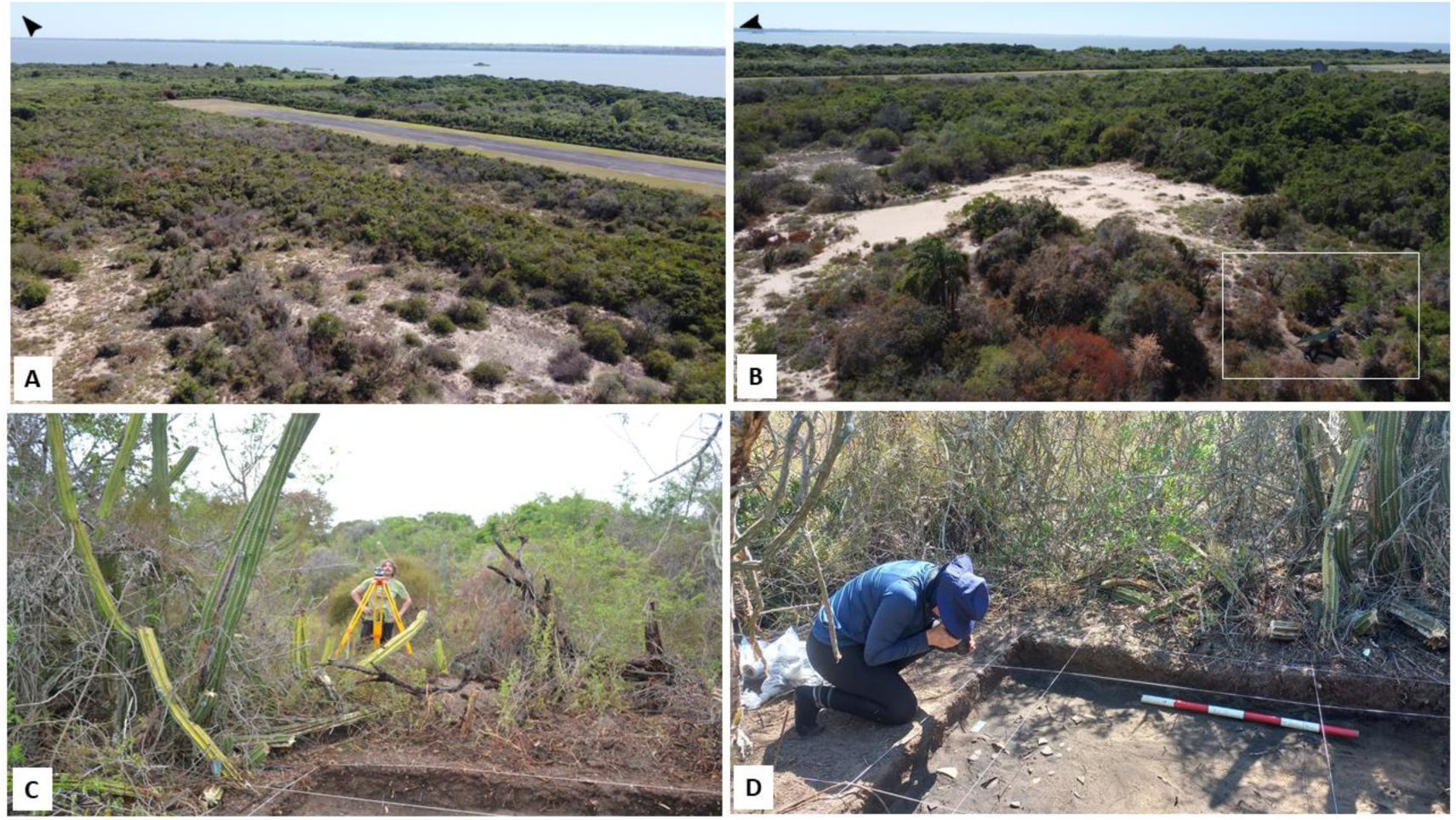
A: Aerial view facing northeast from the excavation area, showing the end of the Arenal Central dune field, the northern tip of the airstrip, and beyond the Del Infierno Channel, the coastline of Martín Chico area. B: Aerial view looking east, with the excavated area from the 2023 season outlined in white. In the background, the airstrip and the Río de la Plata are visible. C: Ground-level view of the forested xerofitic environment surrounding the 2023 excavation area. D: Photographic documentation of the first layer in grid 2. Photograph shows two of the authors of this study (legal disclaimer).

**Figure 6.**
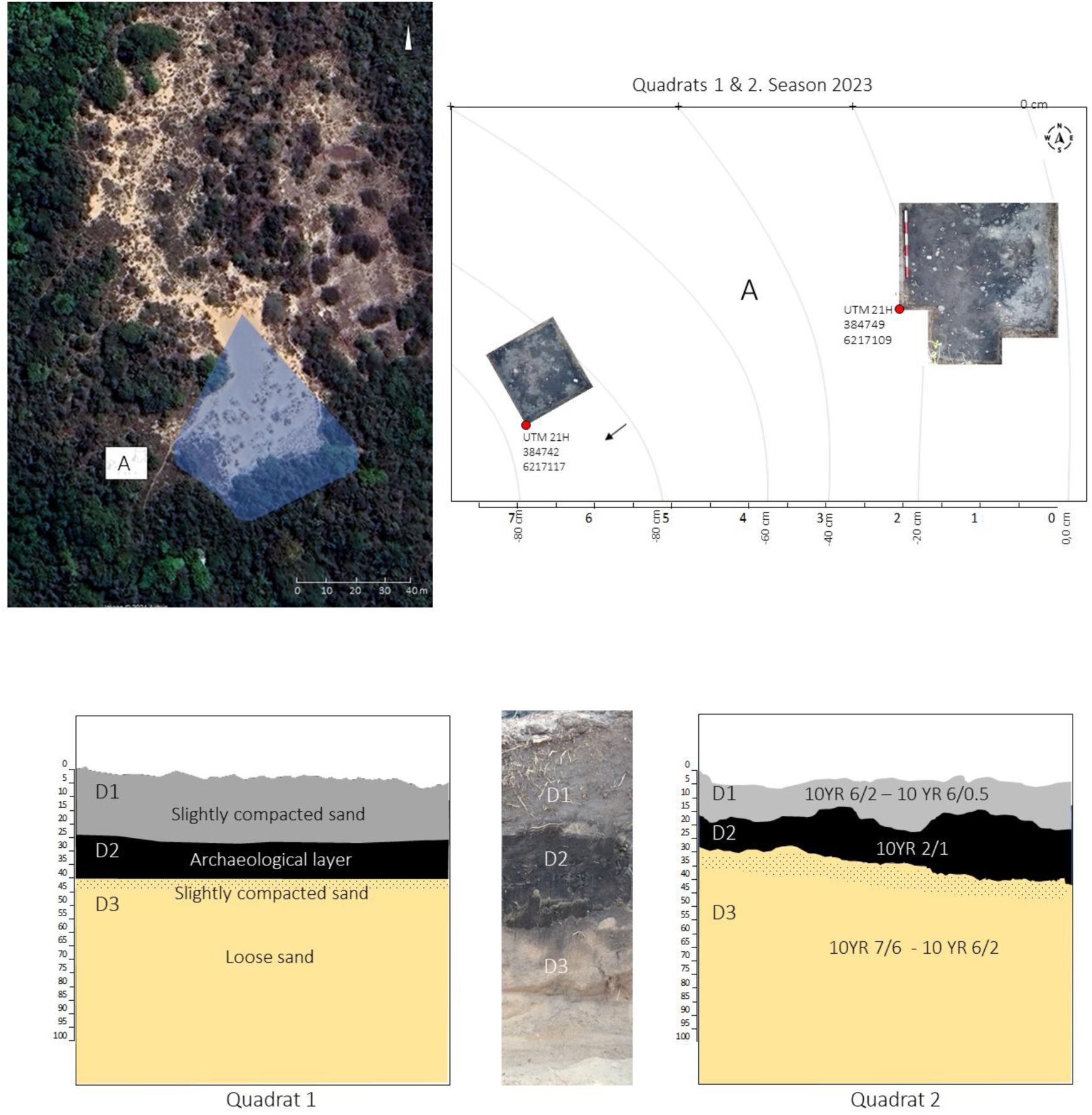
Top: Location of the 2023 excavations (white area marked as “A”) in relation to previous excavations in light blue (approximately located based on Capparelli 2019: 83). Bottom: Stratigraphy of Grids 1 and 2. D1 = slightly compacted loose sand with organic material. D2 = black archaeological layer, slightly compacted. D3 = yellow loose sand, with a somewhat compacted and slightly darkened top.

## 4. Previous archaeological work on Martín García Island

The first archaeological study on the island was conducted by Outes (1917), who published findings of a few Guaraní pottery fragments and human skeletal remains from funerary urns recovered by technicians from the former National Museum of Natural History of Buenos Aires in the Puerto Viejo area (see Figure 4 for the location). In subsequent years, minor interventions were carried out, including one by Vignati (1936). Decades later, Cigliano (1968) excavated “El Arbolito”, imprecisely located somewhere in the northern section of the airstrip, recovering a small collection of Guaraní pottery. Cigliano obtained a radiocarbon date of 405 ± 35 ^14^C years BP (GrN-1456), aligning with the Arenal Central chronology (Section 6.1). Given the island’s size, it is likely that “El Arbolito” and Arenal Central are different areas of a single Guaraní settlement at the northern end of the airstrip.

More recently, Capparelli (2014, 2019) conducted excavations at Arenal Central, covering approximately 80 m² across 17 excavation units. She recovered 2,248 ceramic fragments, a small assemblage of lithic and bone artifacts, and faunal remains. Based on the last comprehensive study of this researcher, ceramic analysis revealed that plain sherds (*n* = 962) and corrugated sherds (*n* = 761) represent 43% and 34% respectively of the total assemblage, followed by fingernail printed (*n* = 242; 10%) and painted (*n* = 283; 13%). Of the 283 painted sherds, all are red monochrome except for six, which exhibit polychromy. Of these six fragments classified as polychrome, Capparelli included five from Cigliano’s previous excavation at the El Arbolito site (Capparelli 2019: 103–104). Thus, only one polychrome fragment was recovered during her excavation.

Reconstructed forms from this collection include corrugated closed-profile pots, likely designed for cooking, small vessels intended for serving food, and large containers probably used for storage, reflecting a wide range of domestic activities. Thin-section analysis of ceramic fragments from this collection revealed that the pastes contained various lithic fragments and large ground sherds as inclusions, which are typical of Guaraní pottery (Pérez et al., 2009, 2018; Capparelli, 2014, 2019; Carbonera & Loponte, 2020; Bertoncello et al., 2024).

The lithic assemblage includes sharp flakes made from cryptocrystalline and silicified limestones, most of which were produced using the bipolar technique. Additionally, slightly modified local pebbles and clasts, shaped through polishing, were identified (Silvestre & Capparelli, 2017; Pérez et al., 2018).

The small faunal assemblage (360 NISP) primarily consists of bony fish species typical of the estuarine environment, along with mammals associated with the island’s riverine-lacustrine habitats and the adjacent plains (Capparelli, 2014, 2019). A more detailed analysis of these findings is provided in Section 6.3. A radiocarbon date obtained from a charcoal sample from Quadrat 3, Level 4, yielded an age of 410 ± 40 ^14^C years BP (LP-2543) (Capparelli, 2014, 2019). This age closely aligns with the one reported by Cigliano (1968). Further details about the site’s chronology are discussed in Section 6.1.

## 5. The 2023 field season

Approximately 7.4 m² were excavated at Arenal Central during the 2003 field season, distributed between Grid 1 (2.4 m²) and Grid 2 (5 m²). The latter was subdivided into smaller sectors (Figure 6). The geographic coordinates of both quadrats are presented in UTM format and reproduced in the sketch in Figure 6, with a margin of error of ± 2 meters. The placement of the grids was determined by the results of 13 test pits conducted in a 100-square-meter area, the condition of the topsoil, and the distribution of vegetation. Since the site is located within a natural preservation area, relatively open spaces were selected for the test pits to minimize vegetation removal, which also influenced the placement, shape, and orientation of the grids. Test pits 6 and 13 yielded positive results, producing a significant quantity of material and exposing dark sediments rich in charcoal fragments. In contrast, the other test pits yielded negative results, with sediments showing only a slight dark staining and few or no archaeological findings. This discontinuous distribution of the archaeological record is typical of Guaraní sites and correlates with the spatial arrangement of residential units within villages (Carbonera, 2014; Brochado, 1984; Goulart, 1987; Métraux, 1948; Rogge, 1996; Schneider et al., 2024b).

The archaeological materials from the grids were uncovered through careful scraping of the sediment, as it is easily removable. The materials were exposed as the excavation proceeded laterally from each one, leaving them uncovered in the process. These exposure levels had a thickness of 2 to 4 cm, depending on the sector and excavation unit, constituting an extraction level. In Grid 1, 85% of the archaeological materials were recovered from the first two extraction levels, while the remaining 15% came from the third and fourth extraction levels. Similarly, in Grid 2, 86% of the materials were retrieved from the first to the third extraction level, with the remainder coming from the fourth and fifth extraction levels. The removed sediment was sieved through a 0.3 mm mesh. All recovered materials are curated at National Institute of Anthropology in Buenos Aires. Materials and methods are described in each section.

## 6. Results

### 6.1. Stratigraphy and chronology

Grid 1 exhibited a slightly consolidated and sandy topsoil with a thickness of 20 to 25 cm. This layer includes modern plant debris, abundant roots, and some redeposited archaeological pottery. Its color, when wet and shaded, ranges between medium gray tones (10YR 6/2 to 10YR 6/0.5). This layer corresponds to an incipiently developed A horizon at the top of Unit D (Ravizza, 1982; see Section 3), referred to here as Layer D1 (Figures 6 and 7). Below Layer D1, between 25 and 40 cm deep, the same sediment from Unit D continues but turns intensely black (10YR 2/1) when wet and shaded, which we identify here as Layer D2. This intense black coloration is attributed to a high concentration of charcoal particles (Figure 7). As the sediment dries, the color lightens slightly. Layer D2 averages 10 cm thick across most of Grid 1, with a maximum thickness of 15 cm. It contains fewer roots than D1 and exhibits a consistent, flat horizontal profile, with only 4 cm of variation across the grid’s extreme points. Despite its limited vertical extent, D2 contains abundant ceramic fragments, faunal remains, lithic artifacts, and both micro- and macro-botanical remains.

**Figure 7.**
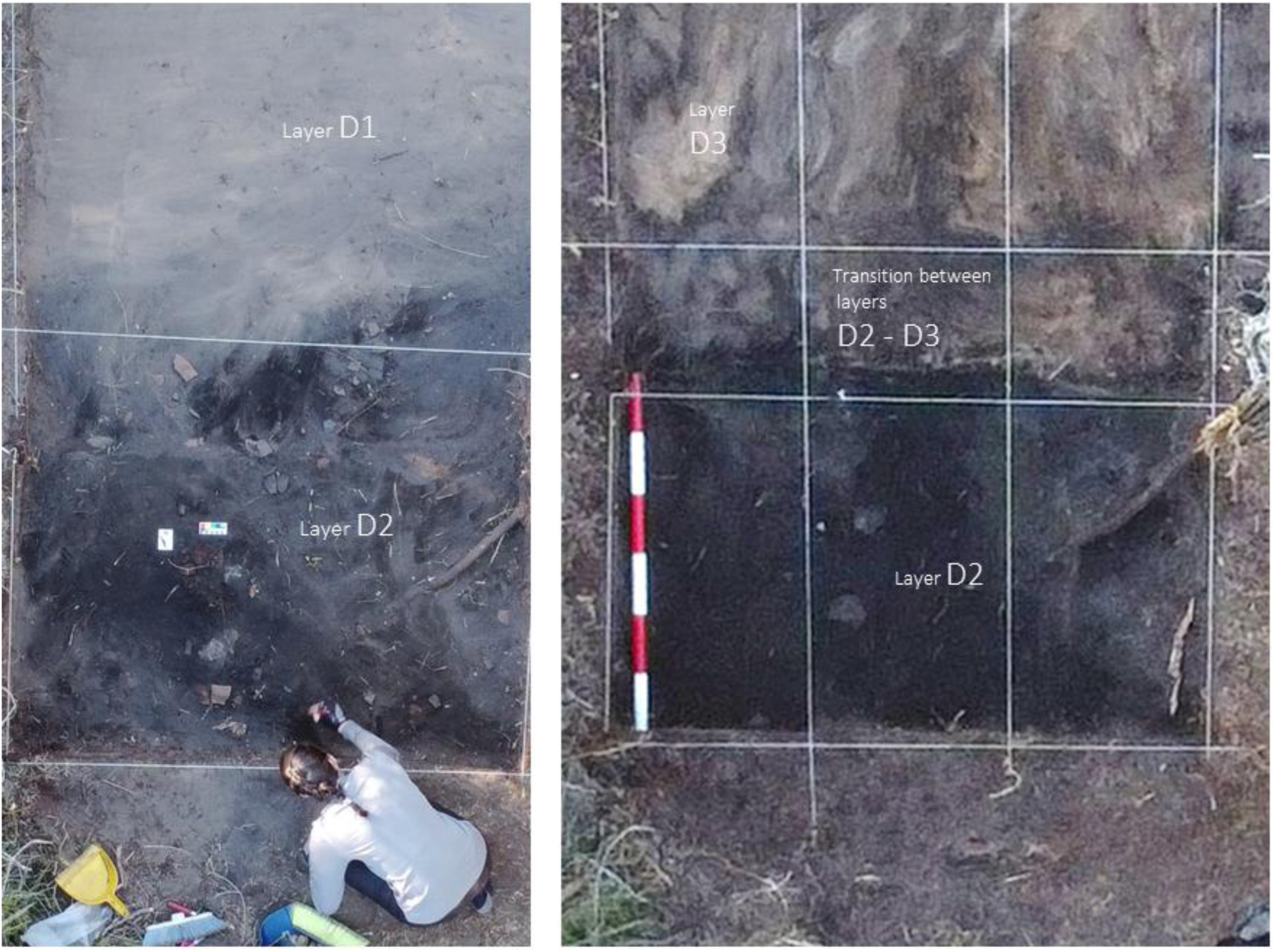
The left image shows Layer D1 at the top of the photograph, with the black-colored archaeological Layer D2 emerging at the bottom. In the right image, Layer D2 is fully exposed at the base of the photograph, followed by the transitional zone between layers D2 and D3 in the middle section of the image, and the upper part of Layer D3 becoming visible near the top edge of the photograph. The scale bar is divided into 20 cm segments. Photograph shows one of the authors of this study (legal disclaimer).

Beneath D2 lies a lighter-colored sand layer with slight tonal variations (10YR 7/6 to 10YR 6/2 when wet and shaded). This layer is also part of Unit D of Ravizza (1982), but for the purposes of the site’s stratigraphy, we identify it here as D3 (Figures 6 and 7). The upper boundary of this layer shows slight compaction and a darker color where it contacts D2. Although D3 occasionally contains isolated archaeological materials, these appear to have descended from D2, making it an archaeologically sterile layer. Approximately 3 to 4 cm below its upper limit, D3 loses consistency and becomes loose yellow sand.

Grid 2 revealed the same stratigraphic sequence as Grid 1 (Figure 6). The thickness of D1 ranged from 11 to 23 cm, while D2 varied between 10 and 15 cm, with small areas extending up to 20 cm. Archaeological materials were primarily concentrated within D2, with a few isolated finds in the top of D3. The greater irregularity of D2 in Grid 2 compared to Grid 1 (Figure 6) likely reflects a higher intensity of phytoturbation, deflation processes, or both.

The distribution and preservation of archaeological materials from both grids demonstrate the good integrity of the D2 archaeological layer, which exhibits a very limited vertical distribution and well-defined upper and lower boundaries. Over 99% of the archaeological materials are confined within this thin D2 layer. Ceramic fragments, lithic artifacts, and bone remains recovered from D2 were predominantly found in horizontal positions, with less than 2% oriented vertically. In Grid 2, a well-defined combustion feature was identified, including large *in situ* fragments of burned logs (Figure 8). This feature extended from the second to the fourth of the five extraction levels. Ceramic fragments and lithic artifacts recovered from Grids 1 and 2 display fresh fracture edges with no signs of rolling or erosion. Several ceramic fragments were successfully refitted, and some lithic flakes from Grid 2 exhibit cortex types that can be linked to one of the cores found within this grid. Bone remains also show angular fractures with sharp edges and no evidence of erosion, corresponding to weathering stage 1 on Behrensmeyer’s (1978) scale, with no signs of scavenging or carnivore marks. Additional indicators of good preservation include very fine, well-preserved fish scales with sharp edges recovered from the sieves, along with both micro- and macro-botanical remains, all in excellent preservation condition.

**Figure 8.**
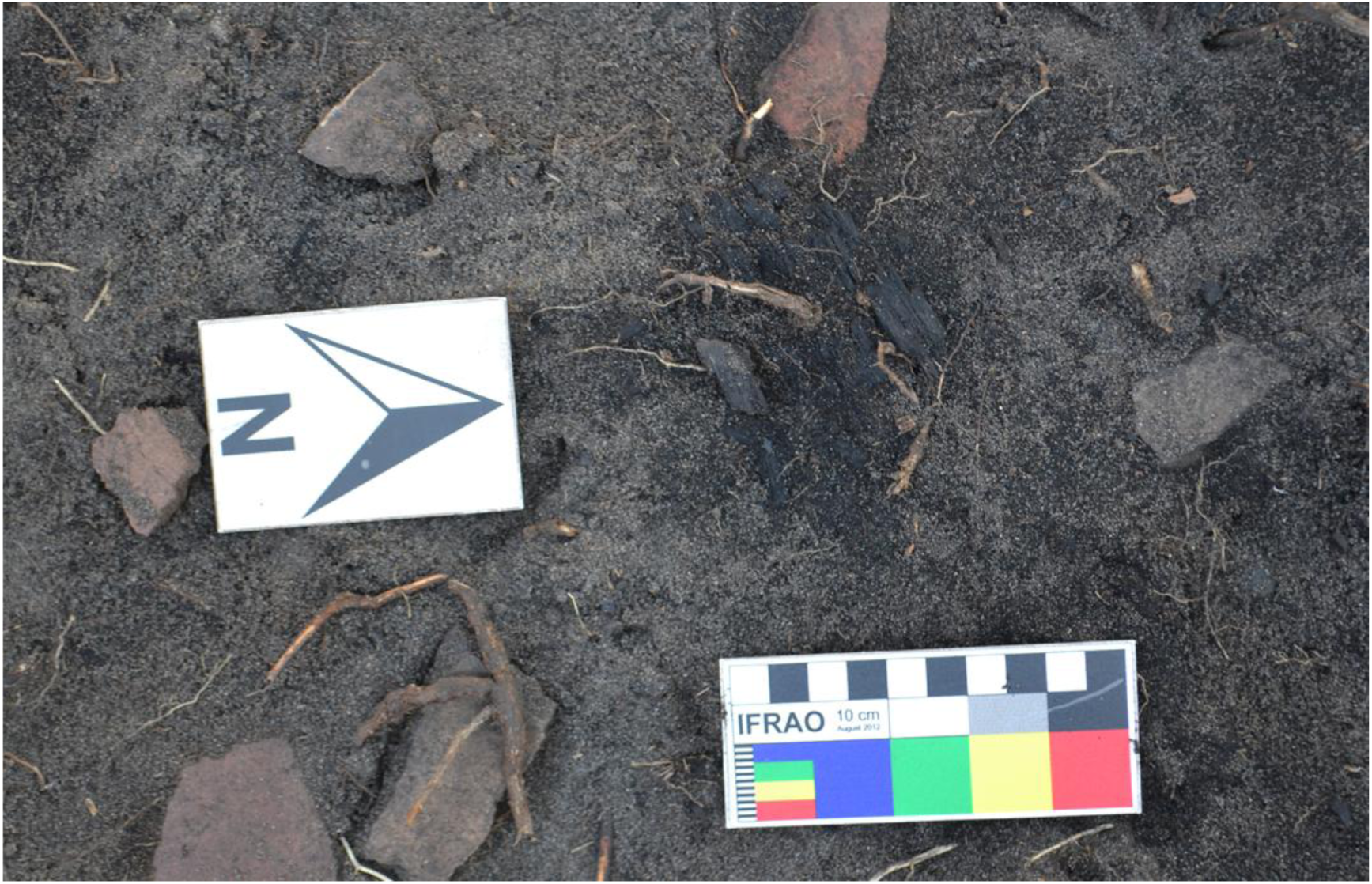
Fragment of a burned log recovered from the combustion structure in Grid 2.

Five AMS dates were obtained from charcoal samples in Grid 1 (Table 1), with calibrated ranges overlapping each other. These include dates reported by Cigliano (1968) and Capparelli (2014, 2019), indicating that Arenal Central was initially occupied around 1450 CE and remained inhabited for several decades (Table 1, Figure 9). The abandonment of the village is further discussed in Section 7.2

**Figure 9.**
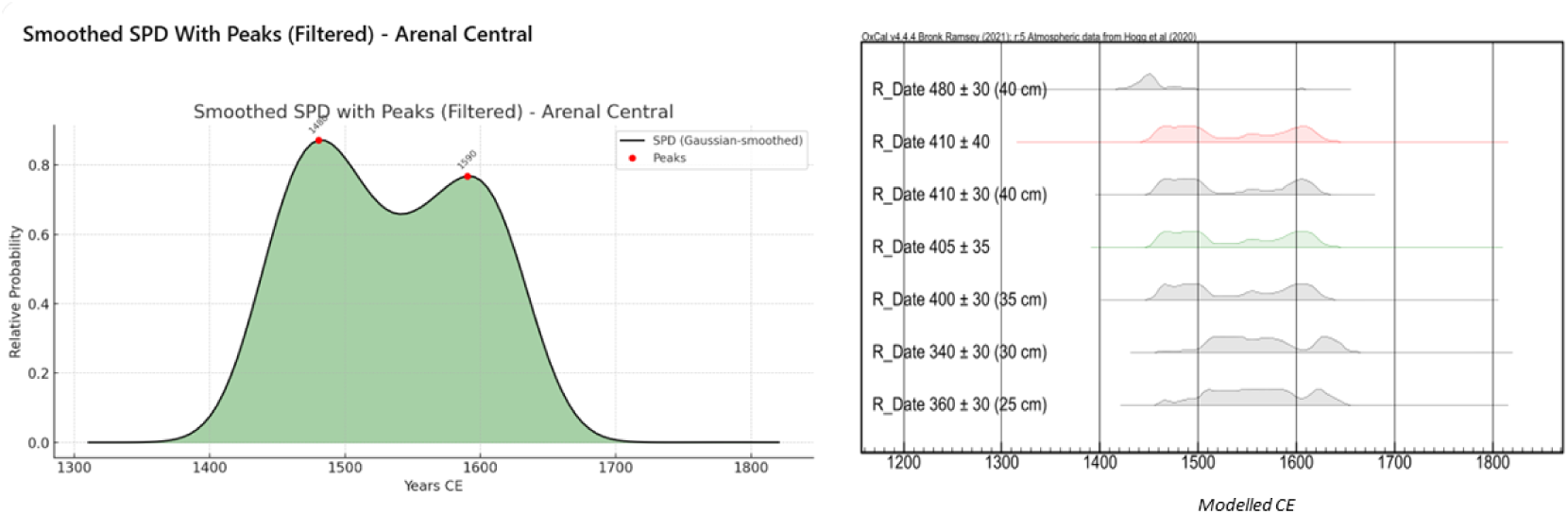
Left: Calibration ranges from Table 1, with Capparelli’s (2014) Level 4 data in red and Cigliano’s (1968) data in green. Right: Summed Probability Distribution (SPD) curve of the dates from Table 1. The curve was smoothed using a Gaussian filter (σ = 10).

**Table 1.** Radiocarbon ages from Arenal Central. All dates have a 95.4% confidence level, except for Beta 662865 (93%) (OxCal, v. 4.4; SHCal-20 calibration curve).

| Site | Quadrat | Depth (cm) | Age $^{14}\text{C}$ AP | Lab. code | Cal CE | Median (CE) | Reference |
| --- | --- | --- | --- | --- | --- | --- | --- |
| Arenal Central | 1 | 25 | $360 \pm 30$ | Beta 662866 | 1483-1644 | 1564 | This study |
| Arenal Central | 1 | 30 | $340 \pm 30$ | Beta 663812 | 1497-1653 | 1575 | This study |
| Arenal Central | 1 | 35 | $400 \pm 30$ | Beta 663813 | 1455-1628 | 1542 | This study |
| Arenal Central | 1 | 40 | $410 \pm 30$ | Beta 663814 | 1451-1627 | 1539 | This study |
| Arenal Central | 1 | 40 | $480 \pm 30$ | Beta 662865 | 1419-1499 | 1459 | This study |
| Arenal Central | 3 | "Level 4" | $410 \pm 40$ | LP-2543 | 1451-1628 | 1540 | Capparelli (2019) |
| El Arbolito | Nd | Nd | $405 \pm 35$ | GrN 5146 | 1452-1628 | 1540 | Cigliano (1968) |

### 6.2. Human Remains

Disarticulated human remains from two adults and one subadult were recovered in Grid 2. These remains were concentrated in the western sector of the grid, distributed radially around the combustion area (Figure 10). The human bones were found mixed with animal bones and covered with charcoal particles, although no direct signs of burning were observed.

**Figure 10.**
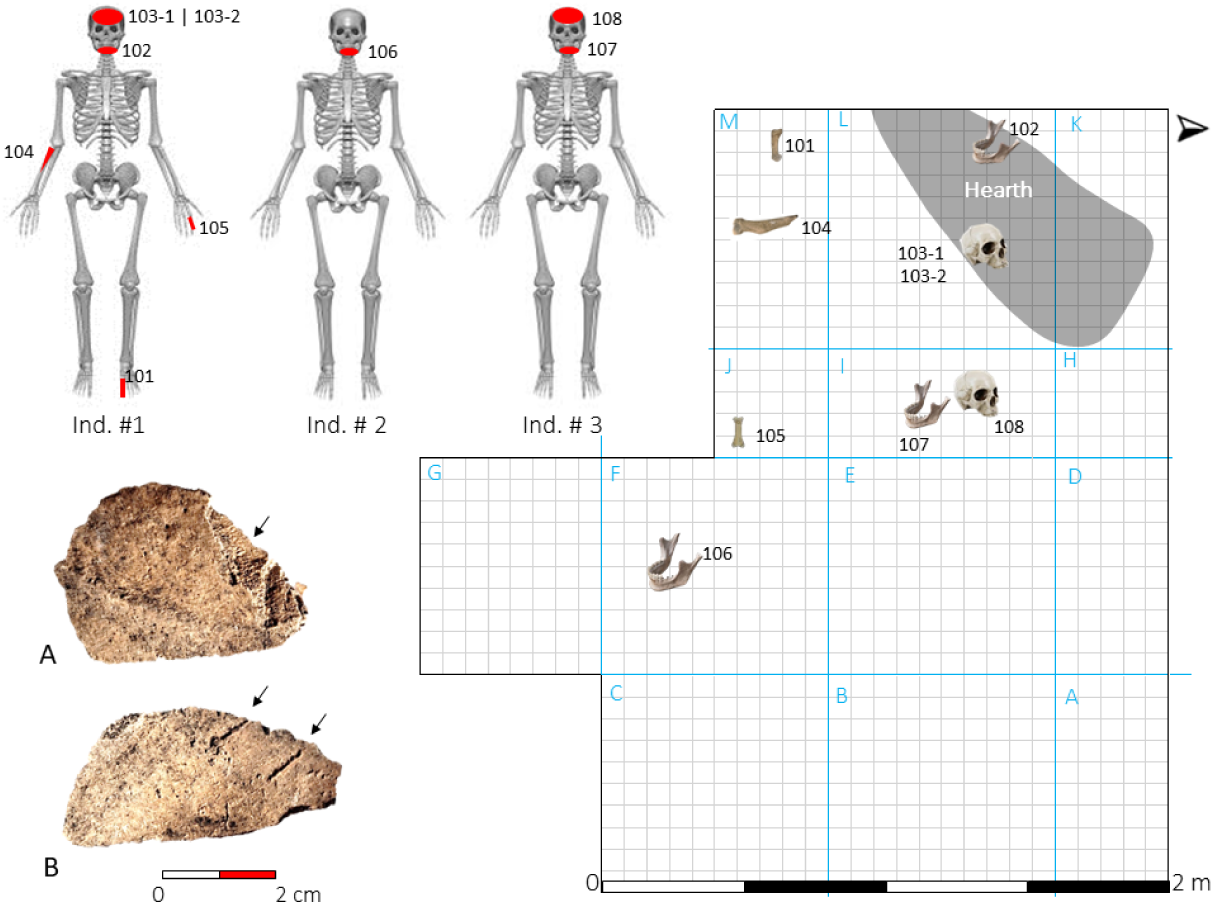
Sketch showing the location of human remains in Grid 2. On the left, skeletal parts represented for each individual (see text regarding the attribution of each fragment). Bottom left: cranial fragment with possible detachment of compact bone tissue while fresh. B: Parallel marks of indeterminate origin (see text).

Individual 1 is represented by cranial bones and several maxillary teeth, identified as 103-1 and 103-2 in Figure 10. The mandible labeled as 102 in the same sketch has been tentatively assigned to this individual, although further studies are needed to confirm this attribution. This mandible exhibits notable weathering, which is atypical compared to other human and faunal remains, suggesting it may have been exposed on the surface of the site for a prolonged period. One of the cranial fragments from this individual display two short, deep parallel marks, which may indicate impact from a blunt object with a slightly convex edge. The groove bottoms are relatively flat and wider than expected for cuts made by a typical lithic flake, suggesting the object was not as sharp as a flake’s edge (Fragment A in Figure 10). Both grooves, along with other areas of this and other cranial fragments, appear to have been affected by chemical dissolution, likely resulting from plant root activity. This affected the edges of both grooves, making them somewhat irregular, though their original design seems to have been strictly straight, as observed at the bottom of both grooves. Another cranial fragment shows evidence of delamination in the compact bone tissue (Fragment B in Figure 10). While no clear percussion point is visible, the nature of the bone loss is consistent with detachment occurring when the bone was still fresh.

Further analyses are ongoing to determine the origin of these modifications, and the results will be presented in future studies. Additionally, two hand phalanges, a metatarsal, and a fragment of the proximal portion of a radius were recovered. These four bones are included under Individual 1 in Figure 10 for illustrative purposes only, as their specific attribution to any individual remains uncertain. Pieces 107 (a fragment of the mandibular ramus) and 108 (two cranial fragments) in the schematic of Figure 10 have been assigned to Individual 2. While Piece 107 undoubtedly belongs to a different adult than Individual 1, the two cranial fragments (108) are tentatively attributed to Individual 2. These fragments feature a significant amount of charcoal adhering to both surfaces. Finally, Individual 3 is a subadult represented by a fragment of a hemimandible (Piece 106 in Figure 10).

The previously identified human remains do not appear to originate from burials, as no evidence of graves was found in the excavated area. Moreover, the representation of the recovered skeletal elements is inconsistent with either disarticulated primary or secondary inhumations, which typically exhibit greater anatomical integrity, with bones that are either unbroken or display dry fractures (e.g., Lothrop, 1932; Loponte & Acosta, 2003-2005; Mazza et al., 2016). Notably, the concentration in Grid 2, consisting of multiple individuals with low skeletal completeness, is similar to that observed in the faunal assemblage, with which the human remains are mixed and arranged radially around the combustion feature. Within this context, what may be processing marks on fragments of the cranium from Individual 1 are also noteworthy. All this evidence suggests that the human remains may have been part of consumption events. In previous excavations, scattered human remains mixed with animal bones were also recovered, with no evidence of disturbed burials identified. One of these bone elements, a phalanx, shows clear evidence of direct exposure to fire (Capparelli, 2019). A similar association of highly fragmented human bones mixed with faunal remains in combustion areas has been documented at Guaraní sites in South Brazil (Rogge, 1996; Mentz Ribeiro, 2008; Spricigo & Carbonera, 2023). These increasingly recurrent occurrences at Guaraní sites require detailed analytical studies to advance their correct interpretation, which will be addressed in future research.

### 6.3. Faunal assemblage

The faunal collection obtained during the 2023 field season includes 307 remains identified at some taxonomic level (Table 2). Approximately 90% of them were recovered between the first and third extraction levels. In previous excavations conducted by Capparelli (2014), 363 bone remains were identified and assigned to some taxonomic category, which are also included in Table 2 as they provide a broader understanding of the recently recovered assemblage. Since the areas excavated by Capparelli are located at a considerable distance from the 2023 excavation (Figure 6), the Minimum Number of Individuals (MNI) values were summed independently; however, some overestimation of the MNI cannot be ruled out.

**Table 2.** Faunal assemblage of Arenal Central. NISP (1) corresponds to the results from the 2023 excavation season, while NISP (2) refers to the remains recovered by Capparelli (2019).

|  | Taxa | NISP (1) | NISP (2) | Total NISP | Total NISP % | Total MNI | Total MNI % |
| --- | --- | --- | --- | --- | --- | --- | --- |
| Mammalia > 10 kg | Mammalia indet. | 83 | 78 | 161 | 24.0 |  |  |
|  | Artiodactyla |  | 1 | 1 | 0.1 |  |  |
|  | *Blastocerus dichotomus* | 2 | 25 | 27 | 4.0 | 2 | 5 |
|  | *Ozotoceros bezoarticus* | 7 | 4 | 11 | 1.6 | 2 | 5 |
|  | *Hydrochoerus hydrochaeris* | 3 | 14 | 17 | 2.5 | 2 | 5 |
| Mamm. < 10 kg | *Myocastor coypus* | 14 | 38 | 52 | 7.8 | 9 | 22 |
|  | *Cavia aperea* | 25 | 6 | 31 | 4.6 | 6 | 15 |
|  | *Holochilus* sp. |  | 1 | 1 | 0.1 | 1 | 2 |
| Aves & Reptiles | *Tupinambis* cf. *merianae* | 1 | 3 | 4 | 0.6 | 2 | 5 |
|  | *Odontophrynus* sp. |  | 3 | 3 | 0.4 | 3 | 7 |
|  | Aves | 4 |  | 4 | 0.6 | 1 | 2 |
|  | *Rhea americana* | 1 |  | 1 | 0.1 | 1 | 2 |
| Fish | Fish | 47 | 148 | 195 | 29.1 |  |  |
|  | Actinopterygii | 114 |  | 114 | 17.0 |  |  |
|  | *Pseudoplatystoma* sp. | 4 | 1 | 5 | 0.7 | 2 | 5 |
|  | *Pimelodus albicans* |  | 5 | 5 | 0.7 | 1 | 2 |
|  | *Prochilodus lineatus* |  | 8 | 8 | 1.2 | 2 | 5 |
|  | *Salminus* cf. *Maxillosus* |  | 1 | 1 | 0.1 | 1 |  |
|  | *Rhamdia* cf. *sapo* |  | 1 | 1 | 0.1 | 1 |  |
|  | *Pimelodus maculatus* |  | 8 | 8 | 1.2 | 2 |  |
|  | *Pterodoras granulosus* | 2 | 18 | 20 | 3.0 | 3 | 7 |
|  | Total | 307 | 363 | 670 | 100 | 41 | 100 |

The 2023 faunal assemblage is exceptionally well-preserved, with approximately 90% of mammal bones from animals weighing more than 5 kg displaying a weathering stage 1 (Behrensmeyer, 1978). The remaining fragments exhibit weathering stages 2–3, most of which were recovered from the first extraction level in contact with Layer D1.

Both Capparelli’s (2014, 2019) assemblage and the 2023 one exhibit a similar pattern, focused on the exploitation of fish and mammals weighing more than 10 kg (Table 2). Fish remains show complete skeletal representation, suggesting that the fish were transported to the site without prior disarticulation. No preference for any particular fish species is observed, a pattern already noted in other Guaraní faunal assemblages (Acosta et al., 2019). Among the exploited mammals, *Blastocerus dichotomus* (marsh deer) stands out as the largest prey (∼up to 150 kg). Its remains, including both axial and appendicular elements, suggest that entire carcasses were transported to the site. However, this species is represented by only a few bones, indicating low completeness in the excavated area (Figure 11). Other identified species include *Hydrochoerus hydrochaeris* (capybara) and *Myocastor coypus* (coypu). The former weighs approximately 30-40 kg, while the latter between 5 and 6 kg. Capybara remains are primarily represented by phalanges and metapodial bones, which also indicates very incomplete anatomical representation in the excavated area, while the coypu shows greater skeletal completeness. These three species could have been hunted on the island, along the riverbanks of the Uruguayan coast, or on the islands at the forefront of the Paraná Delta’s advance.

**Figure 11.**
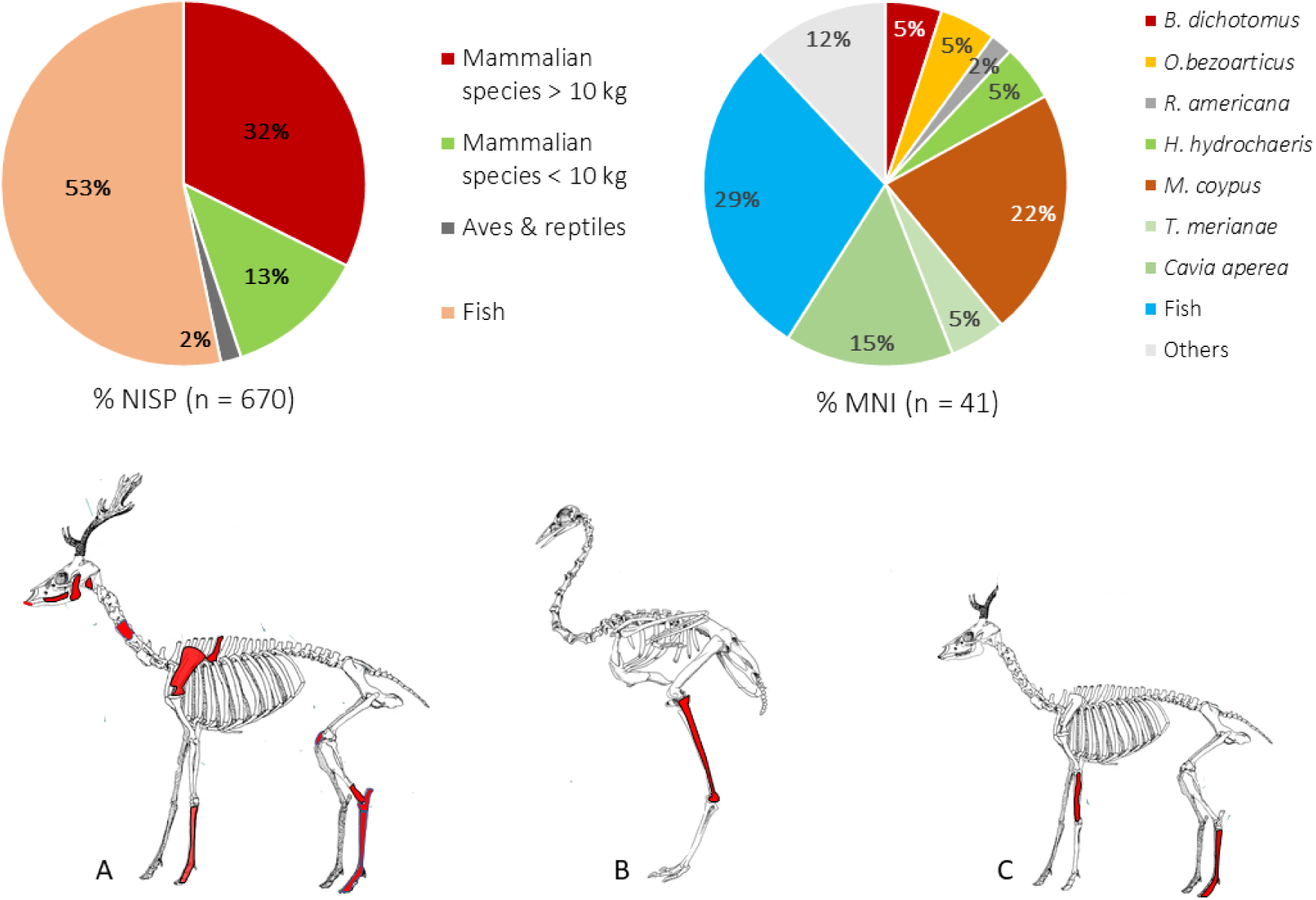
Above: Composition (%) of the faunal assemblage based on total NISP and total MNI values, according to Table 2. Below: anatomical representation of *B. dichotomus* (sketch A), *R. americana* (B), and *O. bezoarticus* (C) based on total NISP from Table 2. Some large bones have been fully colored for better visualization, but only small fragments of them have been recovered (see text).

The faunal collection also includes bones from the forelimbs and hindlimbs of *Ozotoceros bezoarticus* (pampas deer). This small deer (up to 25 kg) could have been hunted in the Uruguayan plains across the Canal del Infierno, less than 4 km away. In the same elevated and well-drained environment, *Rhea americana* (greater rhea) was also available for hunting, from which a proximal fragment of a tibiotarsus was identified (Figure 11).

Some smaller taxa, such as *Cavia aperea* (Brazilian guinea pig) may have entered the archaeological record through natural processes. However, there is no compelling reason to exclude this 300-gram rodent as a potential food resource, given its extensive exploitation by human population across the region during pre-Columbian times (Acosta et al., 2010; Loponte, 2008). Although the bones do not display signs of fresh fractures or fire exposure, a notable concentration of remains of this species was found within the limited excavated area. These remains are highly disarticulated, and the skeletal representation shows low completeness, consistent with patterns observed in other exploited species.

The strategies employed in the exploitation of faunal resources reflect a broad spectrum. The diversity of the 2023 faunal collection, calculated using the NISP values presented in Table 2, yields a diversity index of (Ds) 0.72, which is comparable to that of Capparelli’s collection (Ds = 0.74, NISP). Biodiversity based on the MNI values from Table 2 for the 2023 sample is 0.82, closely aligning with the NISP-derived values. This broad exploitation niche reflects typical Guaraní foraging behavior (Acosta et al., 2019). However, the values from Arenal Central are slightly narrower than those of most other Guaraní assemblages recovered from more northern sites in the subtropical forest. This difference may be attributed to the higher species diversity of the Atlantic Forest compared to the environment of the upper Río de la Plata estuary. Nonetheless, the niche breadth at Arenal Central is broader than that of Arroyo Fredes, the other Guaraní collection studied in the region. The narrower niche at Arroyo Fredes reflects a greater incidence of fish remains (Acosta & Mucciolo, 2010; Musali, 2010a) (Figure 12).

**Figure 12.**
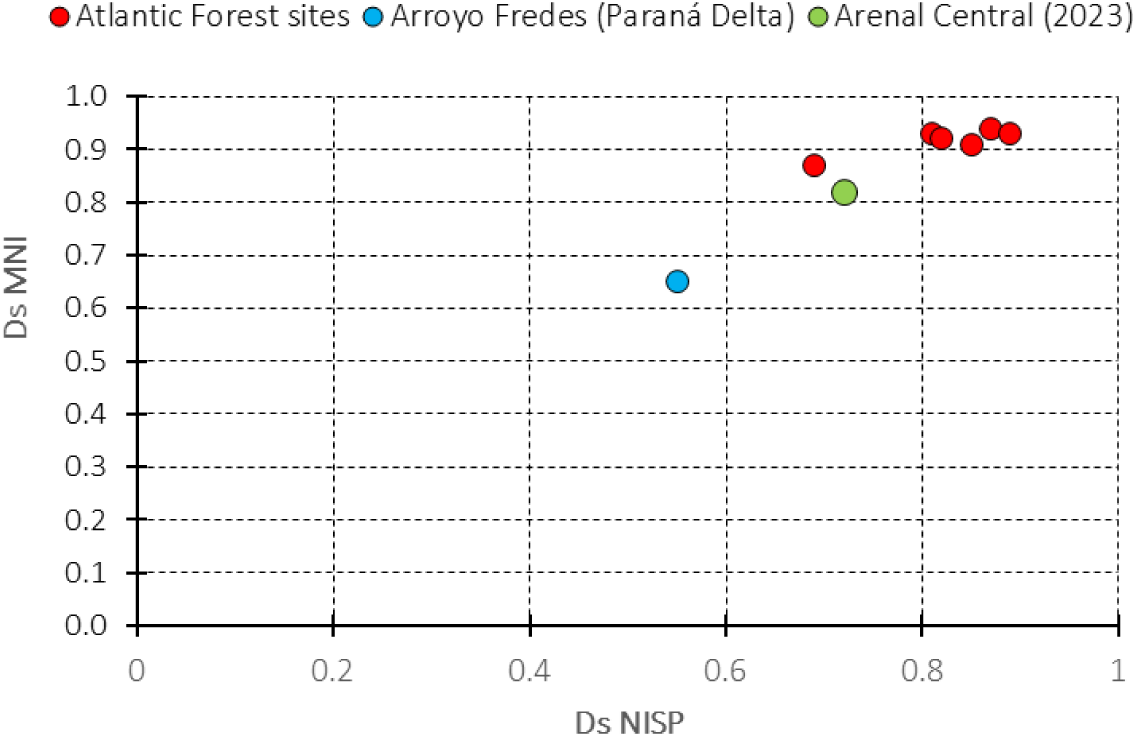
Values of the Simpson (1949) biodiversity index for Guaraní faunal assemblages with adequate published data: Arenal Central, based on the results from the 2023 collection included in Table 2 of this study; Arroyo Fredes, based on data published by Acosta & Mucciolo (2009: Table 1). Additionally, six other sites located in the Atlantic Forest, published by Acosta et al., 2019, were also included. For the calculation of the index, only the number of bone elements (NISP values) and the number of individuals (MNI values) identified at the species level were included, except for fish, which were considered as a single (macro)taxon. Birds were also treated as a single category.

### 6.4. Pottery

A total of 1,230 ceramic fragments larger than 1 cm² were recovered from Grid 2, including 146 rim sherds and 1,084 body and base sherds. Approximately 92% of the fragments were recovered within the first three extraction levels of layer D2, corresponding to only 10 to 12 cm in thickness. The decorative style and technological features observed in the ceramic collection from Arenal Central is entirely Guaraní. The majority of the sherds have either corrugated (∼31%) or plain (∼61%) surfaces (figure 13-A and 13-B). A smaller proportion (∼7%) exhibit painted surfaces (figures 13-C and 13-D), while a minimal fraction (∼1%) display other treatments (figures 13-E to 13-G). Among the 86 recovered painted sherds, nearly 20% are rims.

**figure 13.**
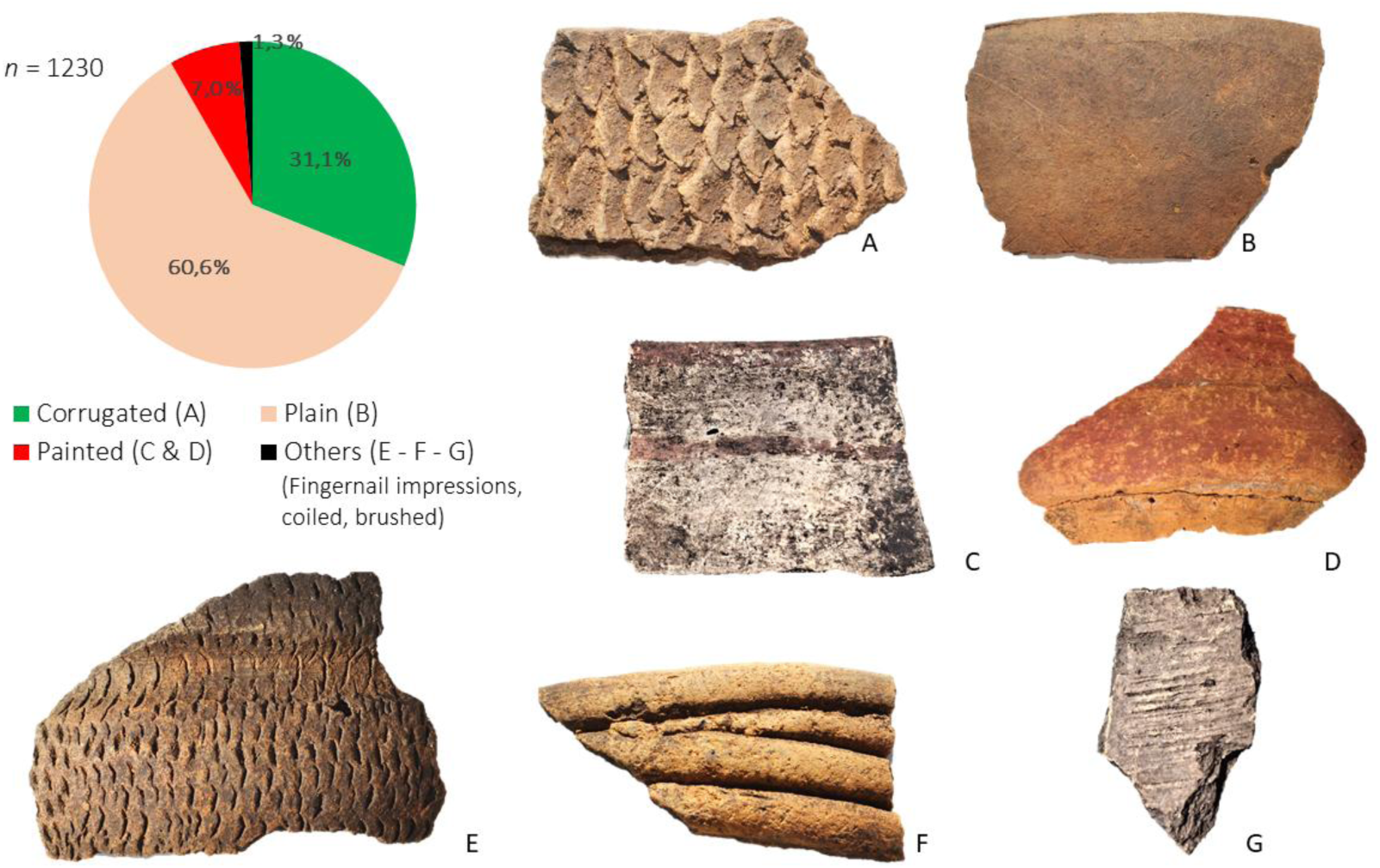
pie graph: surface treatment types of the arenal central ceramic assemblage. a: corrugated surface. b: plain. c: painted (polychrome). d: painted (red monochrome). e: fingernail impressions. f: coiled. g: brushed.

The proportions of the various surface treatment types in the ceramics from Arenal Central fall within the expected range of variation for this type of assemblage, as observed in other collections from residential areas of systematically excavated Guaraní sites. These comparisons, based on analyses conducted using the same criteria (Table 3), include examples from both the immediate region and the upper basin^2^. All these assemblages are characterized by a higher proportion of plain and corrugated pottery, commonly used for domestic purposes (cooking and daily food service). In contrast, in collections made by amateurs or originating from burial areas, the quantity of painted pottery, including polychrome ceramics, significantly increases in frequency (e.g., Lothrop, 1932; Pérez & Alí, 2017: Table 1).

**Table 3.** Surface treatments of ceramic sherds recovered from Guaraní sites located in the upper Paraná Basin and the Paraná Delta–Río de la Plata estuary. References: 1 = Carbonera & Loponte (2025); 2 = Loponte et al. (2022); 3 = Caggiano (1982); 4 = Pérez & Alí (2017); 5 = Capparelli (2019); 6 = This paper. The collection from Paraná Guazú 3 lacks a precise characterization of the type of area that was excavated.

| Collections recovered from Guaraní residential areas in Argentina (rims+bases+body sherds) |  |  |  |  |  |  |  |  |
| --- | --- | --- | --- | --- | --- | --- | --- | --- |
| Sites | Location | Excavated area | Sample (n) | Plain (%) | Corrugated (%) | Painted (%) | Others (%) | Reference |
| Corpus | Upper Paraná River | Domestic house-floor | 4504 | 27.4 | 57.9 | 12.8 | 1.9 | 1 |
| Panambí 3 | Upper Uruguay River | Domestic house-floor | 3850 | 46.8 | 32.3 | 17.3 | 3.5 | 2 |
| Paraná Guazú 3 | Paraná Delta - Río de la Plata estuary | Unknown | 4370 | 36.1 | 18.1 | 40.4 | 5.4 | 3 |
| Arroyo Fredes |  | Domestic house-floor | 2786 | 49.5 | 21.5 | 23.0 | 6.0 | 4 |
| Arenal Central (2019) |  | Domestic house-floor | 2248 | 43.0 | 34.0 | 13.0 | 10.0 | 5 |
| Arenal Central (2023) |  | Domestic house-floor | 1230 | 60.6 | 31.1 | 7.0 | 1.3 | 6 |

Most of the painted fragments are monochromatic, either white or red, as shown in Figure 13-D. The red pigment was applied after the vessels were fired, and appears faint, diluted, and unevenly distributed. In contrast, the white pigment was applied before firing, either as a diluted slip or barbotine. In most of the white fragments, the slip is poorly preserved and, in some cases, has almost completely crackled and eventually worn away.

Polychrome ceramics are exceptionally rare, with only 17 fragments in the entire collection (1.4%). These pieces feature either simple lines or geometric motifs in black or red over a white slip, and less frequently, black lines on a red base. The painted designs are mostly faded and poorly preserved. The degraded state of the painted surfaces contrasts sharply with the excellent condition of plain and corrugated ceramics, as well as other indicators of good archaeological material preservation (see Section 6). The deterioration of the paint may be due to the quality of the pigments used, possibly combined with a solubilization process in porous sediments that promote the rapid infiltration of meteoric water solubilizing salts which attack buried pottery (e.g., Secco et al., 2011). The decorative patterns on the polychrome fragments resemble those documented in Guaraní ceramics from northern regions However, they appear less refined, with broader strokes, reduced geometric precision, and lower complexity (Figure 14) (see also an analysis of some sherds of painted pottery from the island collected by C. Spegazzini and A. Vignati in Torino and Bonomo, 2024). In contrast, the corrugated pottery displays styles and production techniques comparable to those identified in areas of the upper Uruguay basin where proper records and illustrations are available, such as Panambí in the province of Misiones (Sempé & Caggiano, 1995; Loponte et al., 2022), and at several sites along the upper Uruguay River in Brazil (Carbonera & Loponte, 2024; Carbonera et al., 2021).

**Figure 14.**
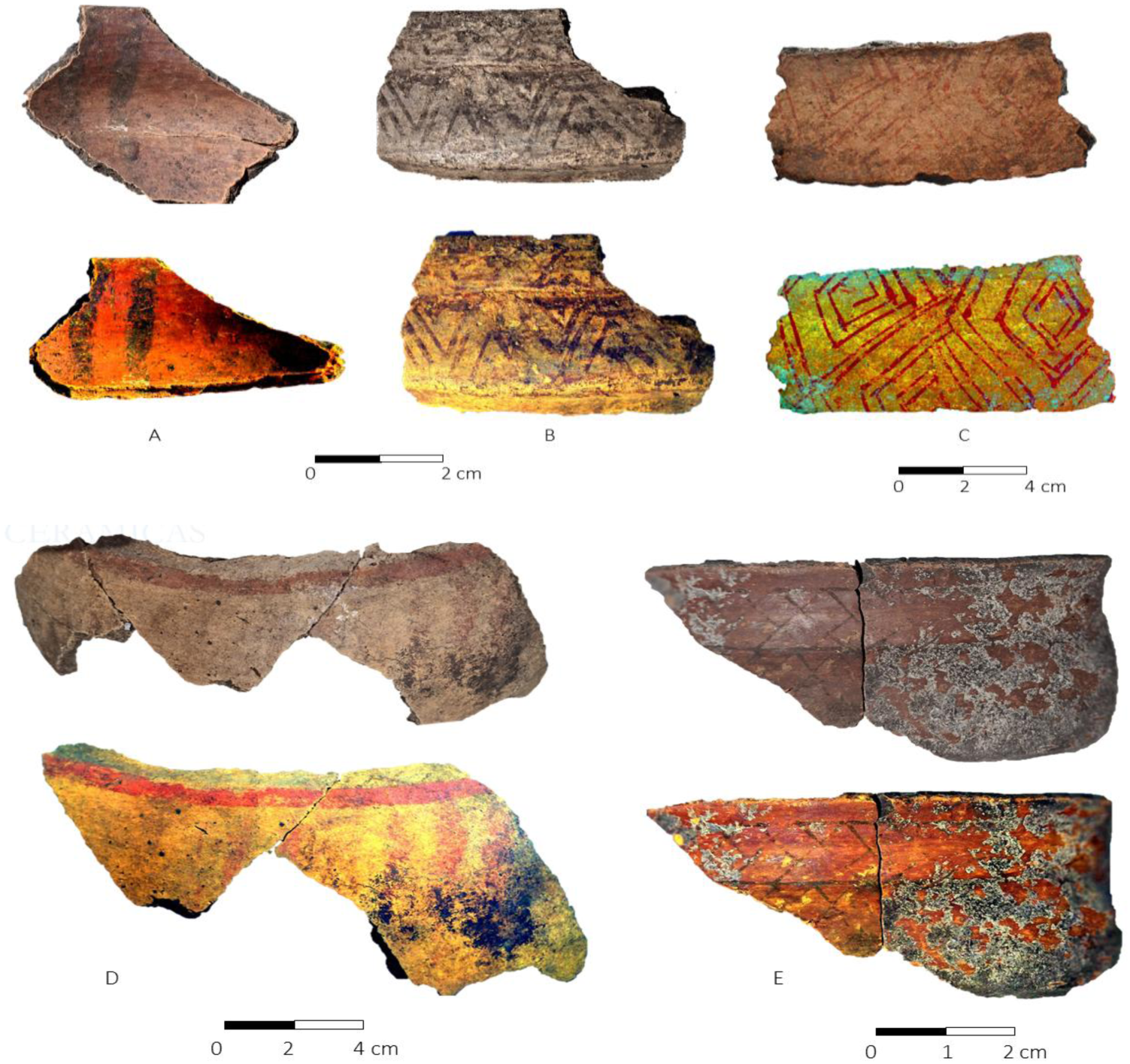
Polychrome ceramic fragments from Arenal Central. Each illustrated fragment is accompanied by its corresponding DStretch-enhanced image. Scales are approximate.

The typological reconstruction of the ceramic assemblage is limited by vessel fragmentation. The reconstruction is further complicated by the low number of fragments per vessel. However, larger fragments and a limited number of refitted sherds allowed us to estimate the shape and size of some of the vessels. Wall thickness also contributed to a better reconstruction of the ceramic assemblage, facilitated by the availability of numerous complete Guaraní pots and reconstructed forms in the literature, notably those compiled by La Salvia & Brochado (1989), Schmitz (1991), Brochado & Monticelli (1994); Prous & Lima (2008), several collaborators included in the latter publication, and additionally by Carbonera and Loponte (2024) specifically focused on the record of the upper Uruguay River also based on complete pots.

One of the reconstructed forms consists of shallow vessels resembling flat plates with slightly concave profiles, featuring corrugated exteriors and smooth interiors, as shown in Figure 15-A. These vessels have an estimated diameter of approximately 40 cm. Due to uncertainty regarding the curvature of the base, they may also correspond to the design depicted in Figure 15-I. Their shape and size suggest they were likely used for serving food. Similar vessels are referred to as *ñaembé* (food-serving plates) by Guaraní communities in early 17th-century Jesuit missions (La Salvia & Brochado, 1989). However, their design also aligns with *nhamopiu* or *ñamipiú* vessels, used as toasters or griddles for preparing cassava tortillas and other semi-solid foods (La Salvia & Brochado, 1989; Brochado & Monticelli, 1994).

**Figure 15.**
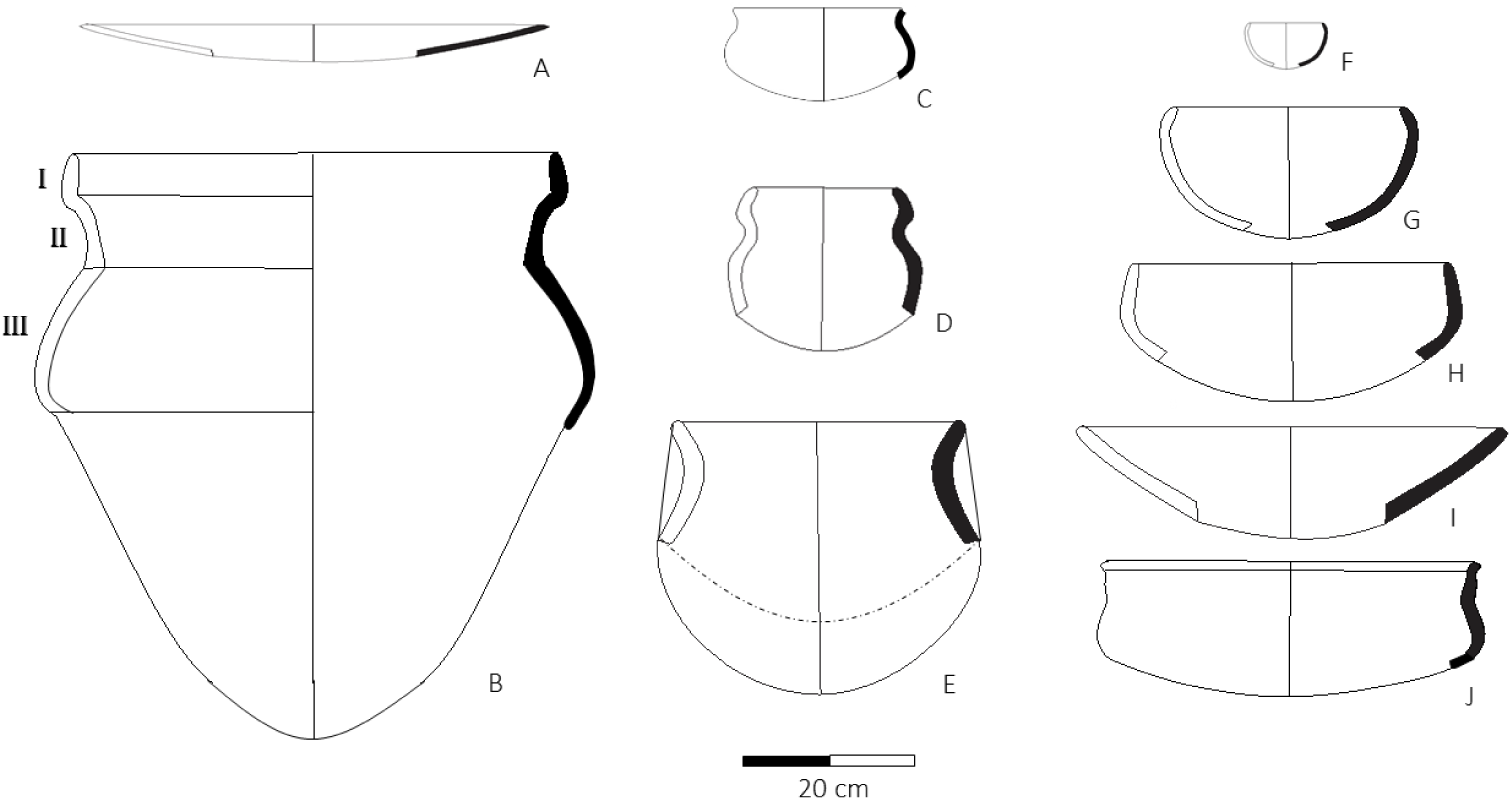
Main typological groups identified at Arenal Central based on the collection from the 2023 season. The thickened area corresponds to fragments that have been identified (see the text above). The scale is approximate

Larger vessels include at least two examples featuring three upper rings, as illustrated in Figure 15-B. Similar containers—some with two or four upper rings—are commonly referred to in the literature as cambuchí (Ambrosetti, 1895; La Salvia & Brochado, 1989; Prous, 2010). Cambuchí vessels are large, robust containers with thick walls easily identifiable. They also display painted side panels, which may be simple and large in size or composed of multiple rings (’stepped shoulders’ in the sense of Prous, 2010: 125), serving as unmistakable identifying features. These fragments cannot be confused with those of smaller and more delicate vessels, such as the cambuchí caguabá or other similarly small containers, nor with the large cooking pots known as yapepó (e.g., La Salvia & Brochado, 1989: 131, 137–139)

One of these vessels is undecorated, while the other displays geometric red patterns painted over a white background across the three rings (upper, middle, and lower), a distinguishing feature of these pots. A fragment from the lower ring of the latter is illustrated in Figure 14-C. The fragmentation of these pots generates a large number of painted sherds from the upper half of the containers and plain fragments from the lower half. Vessels like these were probably used for preparing and storing fermented beverages and other liquids (Ambrosetti, 1895; La Salvia & Brochado, 1989; Brochado & Monticelli, 1994).

Medium-sized vessels include pots with slightly inverted rims, like the one shown in Figure 15-E, although some also feature straight or even slightly closed rims (same figure). Since no fragments connect the rims to the bases, these vessels could have had an abrupt carination, defining a pot with a short profile, or a very gentle carination resulting in a well-developed base corresponding to a pot with a taller profile and deeper base. These options are illustrated in Figure 15-E. Historically, these vessels were used for cooking food and are referred to as *yapepó* by ethnographic Guaraní communities, with numerous variations depending on the specific form (Ambrosetti, 1895; La Salvia & Brochado, 1989; Brochado & Monticelli, 1999). Similarly, the Asurini people along the Xingú River, also part of the Tupi-Guaraní linguistic family, use the term *japepá* for similar vessels (Silva, 2010). Most of these cooking pots exhibit corrugated walls and often contain abundant charcoal particles on their exteriors, indicating use over fire.

Another type of pot, with straight or slightly inverted profiles, corresponds to the vessel depicted in Figure 15-G, along with smaller forms like the one shown in Figure 15-F. These vessels bear ethnographic names such as *ñaetá* or *caguabá*, depending on subtle variations in shape and size (La Salvia & Brochado, 1989). Both terms refer to containers used for serving food and beverages. These vessels generally have thinner walls than the cooking pots described earlier and show no evidence of fire exposure, further supporting their use as food-serving containers. While most of them feature plain surfaces, some smaller variants exhibit corrugation.

Three other vessel types, also typically Guaraní, were identified and are illustrated in Figures 15-D, 15-H, and 15-J. The first type consists of thin-walled vessels with a waist in the middle or upper third of the body, which never show signs of fire exposure. This suggests that they were not used for cooking but rather for storing and serving liquids. A fragment of this type is illustrated in Figure 13-E. Similar vessels, known as *yawi* by the Asurini, are used for transporting and storing liquids (Silva, 2010). Ethnographic Guaraní also used small vessels with a narrow waist in the middle or upper third of the body, known as *cambuchí yaruquaí*, for storing or drinking liquids (La Salvia & Brochado, 1989). The remaining two types are low-walled vessels with slightly convex bases. Their walls are smooth and often painted up to the carina on the exterior. The paint typically consists of a white slip as the base, decorated with red or black geometric patterns or a simple red line parallel to the rim. Some also feature additional external geometric motifs, as seen in fragments 13-C and 14-B. A few fragments of vessels of this type display red paint as the base color, as observed in fragment 13-D. The interiors of these vessels are usually painted monochrome red, occasionally incorporating geometric designs. At Arenal Central, these interior motifs are simple, such as those in Figure 14-A, where the black lines appear to have been applied by finger-painting. The open, low-walled design suggests that these vessels were primarily used for serving food and beverages. Ethnographic parallels include *cambuchí caguabá* vessels, traditionally used for consuming fermented drinks (La Salvia & Brochado, 1989).

These are ethnographic functional categories and should be regarded as such. Nonetheless, they exhibit substantial temporal persistence, as they have remained consistent from the 17th century to the present, and are found across different ethnographic groups within the Tupi-Guarani language family (Ambrosetti, 1895; Brochado & Monticelli, 1994; La Salvia & Brochado, 1979; Noelli et al., 2018; Prous, 2010; Silva, 2010).

Within the ceramic assemblage, other vessel fragments have been identified that appear to be subtle variations of those previously described, as well as others with different morphologies. The incomplete nature of these fragments prevents a confident reconstruction of their profiles. Therefore, the typological reconstruction presented should be considered a minimum estimate of the typological diversity. In Figure 15, we present reconstructions of the identified vessel types. Following a common convention in archaeological publications worldwide, certain sections of the profiles have been thickened to indicate the fragments from the 2023 collection that enable the reconstruction of the original vessel shapes (e.g., Chirikure et al., 2013; Navarro et al., 2022; Stronach et al., 2019, among hundreds of published studies).

### 6.5. Bone Artifacts

Five bone artifacts were recovered during the 2023 excavation season, four of which are illustrated in Figure 16. Among them is a projectile point crafted from the compact tissue of a long bone from a big mammal (Figure 16-A). This point is not Guaraní but exhibits all the technological and stylistic traits typical of bone points from hunter-gatherer sites located on the right bank of the Paraná River and within the Paraná Delta (Loponte, 2008; Buc, 2012). Its distinctive features include an elongated, extremely thin blade, sharp barbs that likely become straight through successive fractures, a short quadrangular stem, and a groove along the center of the blade, probably designed to enhance the projectile’s penetration capability. Two additional artifacts, likely used as projectile tips, were also recovered. One is a small fragment severely damaged by fire, while the other is made from the diaphysis of a medium-sized mammal, such as a coypu. Bone points like these occasionally appear in Guaraní contexts (Figure 16-B).

**Figure 16.**
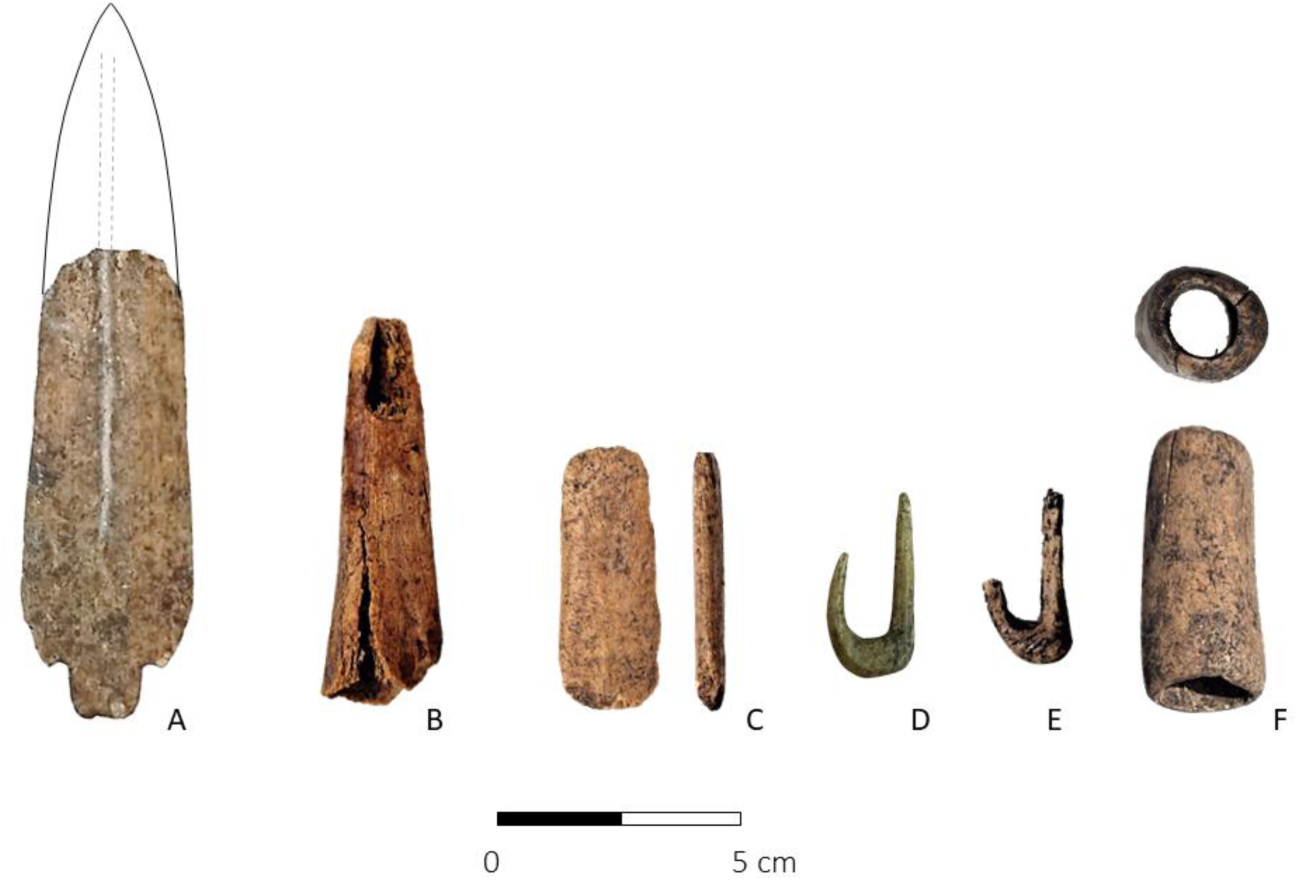
Bone artifacts from Arenal Central. A: Stemmed projectile point with (broken) barbs. B: Hollowed point. C: Polished bone splinter. D: Fishhook (Season 2023). E: Fishhook (Capparelli, 2019). F: Fragment of a diaphysis, cut at both ends (Capparelli, 2019). The scale is approximate.

The fourth artifact (Figure 16-C) is a polished fragment of compact bone resembling a wedge of unknown use. The final artifact is a small fishhook (Figure 16-D), similar to another one recovered during the previous excavation of the site (Capparelli, 2019) (Figure 16-E). Although still underrepresented in the archaeological record, such artifacts are becoming increasingly common in Guaraní contexts. Comparable fishhooks have been identified at the Leandro Meier site (Upper Uruguay River), excavated by the authors (unpublished), and at Aldeia Ribanceira (Atlantic coast) (Demathé, 2023). Capparelli (2014, 2019) also recovered a cut and polished diaphysis fragment (Figure 16-F), whose function remains uncertain but could have been used as an inhalator, given its similarity to ethnographic pieces designed for that purpose (e.g., Ostapkowicz, 2020).

### 6.6. Lithic Artifacts

The lithic assemblage recovered during the 2023 season consists of 24 artifacts, including four axe heads. Three of them appear to have fractured after use or during manufacture, while the largest one was likely still in the process of production (Figure 17). All of these artifacts seem to have been made from local rocks of the Martín García Complex. However, petrographic analysis will be necessary to accurately determine the specific lithologies.

**Figure 17.**
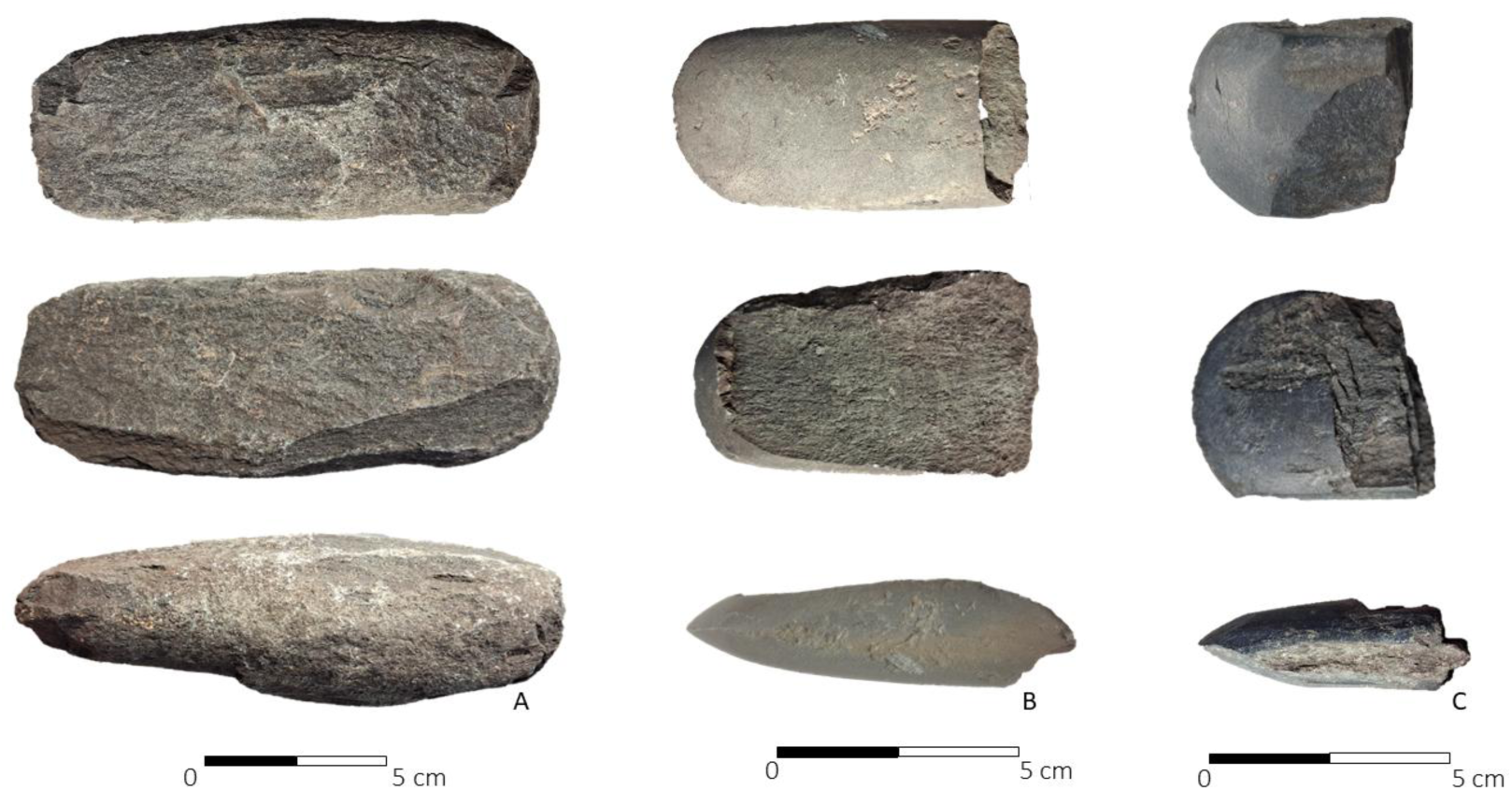
A: Unfinished axe head. B and C, fragments of broken axe heads recovered from Grid 2. The scales are approximate.

A second group of artifacts consists of five river pebbles made of microcrystalline or cryptocrystalline rocks that are typical of the Uruguay River valley. Three of them show evidence of flake extraction using bipolar knapping techniques (Figure 18-B). A fourth pebble has a single percussion mark, while the fifth remains unmodified.

**Figure 18.**
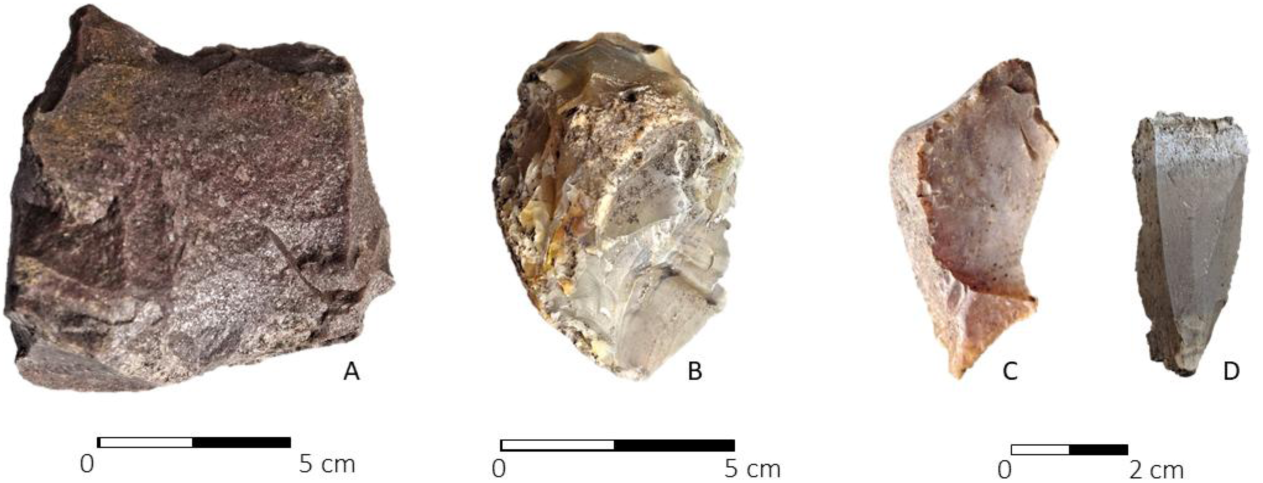
A: Silicified sandstone core. B: Microcrystalline rock (fluvial pebble). C and D: Flakes from microcrystalline rocks. The scales are approximate.

Another group includes 11 small flakes with natural edges, each no longer than 6 cm (Figure 18-D), which appear to have been extracted from the previously mentioned pebbles. A silicified sandstone core was also recovered, showing evidence of multidirectional flake extraction through direct percussion (Figure 18-A), along with two fragments of local igneous-metamorphic rocks, both bearing signs of polishing. One of the fragments contains traces of ochre on its polished surface, while the other may have been used as a mortar. In addition, four small pieces of pale red ochre, each no larger than 3 cm, were recovered.

## 7. Discussion

### 7.1. Site structure and function

Arenal Central represents a Guaraní occupation on sedimentary Unit D, resulting in a thin archaeological layer that reflects various domestic activities, including the preparation, consumption, and storage of food and beverages, as well as the production, use, and disposal (intentional or not) of lithic and bone artifacts. This pattern aligns with what is typically described as a ‘house floor’ record (LaMotta & Schiffer, 1999), and similar to several Guaraní sites excavated is South Brazil and Argentina.

This domestic area also contains a combustion feature that appears to structure part of the assemblage in Grid 2. This feature likely remained active for an extended period, undergoing periodic maintenance and cleaning. The ceramic collection from this house floor record consists of numerous vessels, each represented by only a few small fragments (see Section 6.4). These characteristics suggest a formation process involving repeated cycles of use, breakage, cleaning, and replacement of pots over time, resulting in the observed pattern. A similar sequence— preparation, consumption, disposal, and cleaning—explains the formation of the faunal assemblage, which primarily consists of small bones. The human remains found in Grid 2 might also represent food-related disposal, given their spatial arrangement, completeness, and fragmentation.

The small crystalline rock cores described in the previous section were common within these domestic areas, where flakes were extracted as needed for household tasks. The identification of at least one axe head that clearly appears to be in the process of manufacture suggests that these tools, or at least some of them, were produced within the domestic sphere.

Findings from previous studies (Capparelli, 2014, 2019; see also Table 1) indicate that the occupied area at Arenal Central is considerably extensive, consistent with expectations for a Guaraní residential base (e.g., Métraux, 1948; Schneider et al. 2024b). The discontinuous distribution of the archaeological record, limited to specific areas of the terrain that align with sediment enriched with charcoal particles, reflects a typical pattern observed in Guaraní villages, where residential units were arranged around a central plaza (e.g., Métraux, 1948; Schneider et al., 2024b). The important point here is to identify the scale of the investigations conducted both at this site and at other Guaraní sites, where excavations are usually limited in extent and only represent specific sectors of the sites. Ongoing studies will provide additional insights into site variability.

### 7.2. Chronology

While the Spanish crown discovered America in 1492 CE, the Río de la Plata became part of Europe’s known geography in 1516 CE, when a Spanish fleet led by Juan Díaz de Solís first navigated the estuary (Madero, 1939) ^3^. The earliest radiocarbon date from Arenal Central (Beta 662865; Table 1) falls between 1419 and 1499 CE (93%), leaving only a minimal probability (1.4%) of overlapping with the historical period (Figure 19). However, the remaining radiocarbon dates range from the pre-European period to approximately 1600 CE, well into the historical period. This broad chronological range results from a plateau and a minor reversal in the calibration curve, spanning approximately 1460 to 1640 CE (Figure 19), which prevents more precise dating. This issue is similar to the ‘Ib’ disturbance in the Northern Hemisphere, now dated to approximately 1540–1640 CE (cf. Taylor et al., 1996; Hogg et al., 2020). This challenge limits high-resolution dating for this critical period, which coincides with the onset of early historical times. It is, therefore, possible that the occupation of Arenal Central overlapped with European exploration of the Río de la Plata and the establishment of the earliest Spanish settlements in the region.

**Figure 19.**
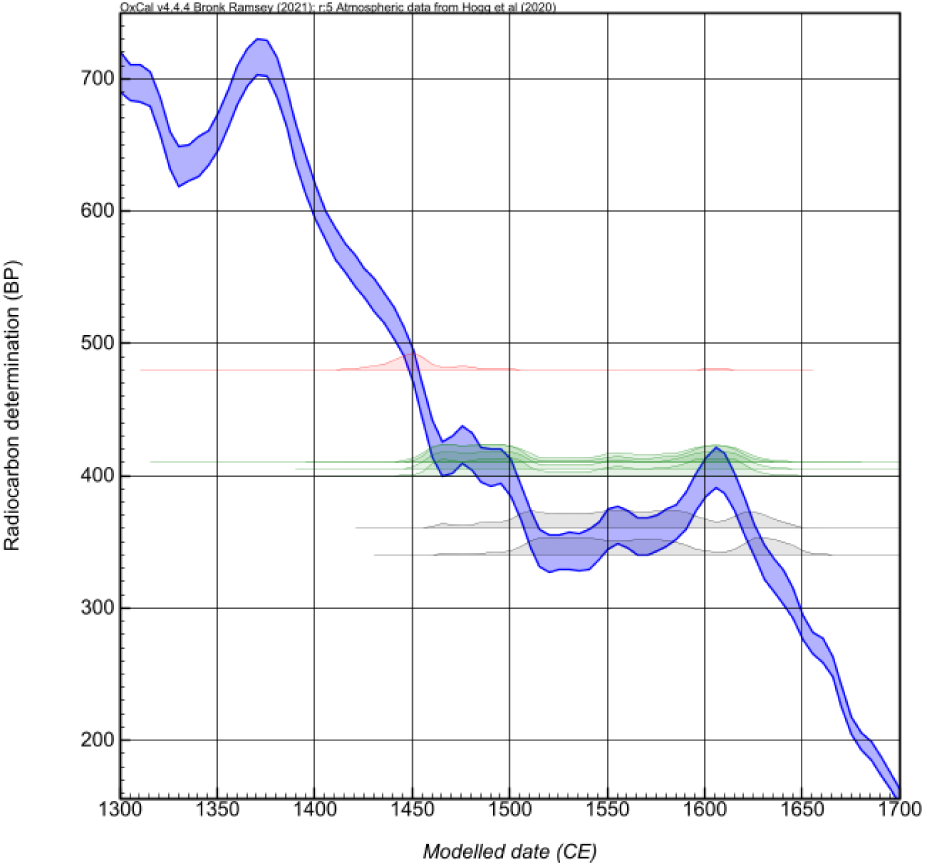
Distribution of the calibrated ranges of the dates from Table 1. Note the presence of a plateau and a slight reversal in the calibration curve where the dates from Arenal Central – El Arbolito are located, except for Beta 662865 (in red), which falls within a well-sloped area of the curve, with most of its range (93%) corresponding to precolonial times. The dates in green (Beta 663813, Beta 663814, LP-2543, and GrN 5146) also have their lower ranges in precolonial times (⁓1450 CE), while the dates in gray (Beta 662866 and Beta 663812) exhibit lower ranges closer to the historical period (1483 CE and 1497 CE, respectively; see Table 1).

Some historical events shed light on the timing of Arenal Central’s depopulation, offering an opportunity to discuss the site’s abandonment in depth. During the first European expedition to the Río de la Plata in 1516 CE, a crew member of this fleet, Martín García, reportedly died of natural causes and was buried on the island, according to Fernández de Oviedo y Valdés ([ca. 1526–1557] 1945). If accurate, this account suggests that the island was already uninhabited at the time. Subsequent events further support the absence of a local population. After this stopover, Solís attempted to ‘capture a man to take to Castile’ (Herrera y Tordesillas [1601–1615] 1944–1945), implying that the small island had no inhabitants. While it is possible that indigenous groups employed a strategy of ‘avoiding contact’ with Europeans, it would have been nearly impossible to hide an entire village the size of a Guaraní settlement—or even a smaller one—on an island as small as Martín García. For this reason, the lack of mention of an abandoned village suggests that it had already been deserted and overgrown with vegetation some time earlier.

To achieve their objective of capturing a man, Solís and his crew immediately landed on the Uruguayan coast, ‘near Martín García Island’ (Herrera y Tordesillas [1601–1615] 1944–1945), possibly at Martín Chico or a nearby northern area. Here, the Spaniards observed indigenous people signaling to them from the beach. According to accounts from the remaining crew members aboard the anchored ships, Solís and several of his men were killed upon landing, then roasted and eaten, while the helpless crew watched from a distance, unable to intervene. While interpretations of this event in local historiography vary, the plausibility of the attack as described is supported by the consistency of the crew member testimonies, particularly if Guaraní groups were responsible for the deaths (cf. Herrera y Tordesillas [1601–1615] 1944–1945).

Regardless of the precise sequence of events described above, other evidence suggests that the island remained uninhabited in the years following Solís’s expedition. In 1528 CE, Luis Ramírez, a member of Sebastián Gaboto’s expedition to the Río de la Plata, visited an ‘island in the middle of the river.’ Accompanied by Indigenous guides, Ramírez stayed on the island for several days and described it as uninhabited and lacking food resources (Ramírez, 1528, in Madero, 1939). While Madero (1939) and Outes (1917) argue that this island corresponds to Martín García, some doubts remain regarding this identification, especially considering that the Río de la Plata estuary had a different configuration than it does today. Nevertheless, Ramírez’s account is significant in broader terms, as it describes the entire area without mentioning the existence of nearby Indigenous villages.

Three years later, in 1531 CE, Pero Lopes de Souza ([1530–1532] 1952) likely visited the island, naming it ‘Santa Ana,’ and likewise noted the absence of inhabitants. Shortly thereafter, the first foundation of Buenos Aires took place in 1536 CE, directly across from Martín García. Buenos Aires was abandoned a few years later, in 1541 CE, when Domingo Martínez de Irala depopulated the small town. In a document addressed to future settlers, Martínez de Irala ([1541] in Zeballos, 1898) described the island (“Martín García” in the original text) as a suitable location for raising pigs. The letter’s subject, the absence of any mention of Indigenous people on the island, and its proposed use for pig farming suggest that the island was uninhabited, consistent with previous accounts.

Several years later, in 1573 CE, the army led by Juan Ortíz de Zárate was defeated by Indigenous groups in the Battle of San Gabriel on the Uruguayan plains. The survivors sought refuge on the island (“Martín García” in the original text) for several days, which was again described as uninhabited (Rela, 2003).

Finally, no European materials or exotic fauna were found during excavations at Arenal Central (Capparelli, 2014, 2019; this study). Based on these historical references and the absence of European materials, it is likely that the Guaraní population abandoned the island shortly before the onset of the historical period, possibly around 1516 CE, leaving no opportunity for exchange processes. However, minor or intermittent reoccupations cannot be entirely ruled out until ∼1600 CE, when the Guaraní archaeological signal disappears as an independent unit throughout the entire area (Section 7.5).

### 7.3. Geographic location of Arenal Central during the Guaraní occupation

Various estimates based on historical cartography from 1750 CE to the present indicate that the progradation rate of the advancing front of the Paraná Delta averages between approximately 30 and 100 meters per year, depending on the specific sector of the Delta and the time period considered (Pittau et al., 2005; Medina and Codignotto, 2013). This implies that most Guaraní sites in the current Paraná Delta were established on sandy islands within the Río de la Plata estuary, located several kilometers away from the advancing Delta front. The key question now is whether it is possible to estimate how far these settlements were from the advancing front.

Using the most conservative average progradation rates reported by Pittau et al. (2005) and Medina and Codignotto (2013), the approximate location of the advancing front in 1500 CE can be estimated, as shown in Figure 20, assuming an isometric advance equivalent to the rates observed after 1750 CE. This estimate not only relies on the most conservative progradation values, but also adopts a cautious approach to inferring the position of the Delta front— particularly through comparison with the overlapping maps of Bellin (ca. 1759 CE) and Juan de la Cruz Cano y Olmedilla (1775 CE), which provide highly detailed information on the historical position of the advancing front, including the distribution of estuarine islands and the course of right-bank rivers such as the Luján River, which is key to reconstructing the former extent of the Delta’s leading edge.

**Figure 20.**
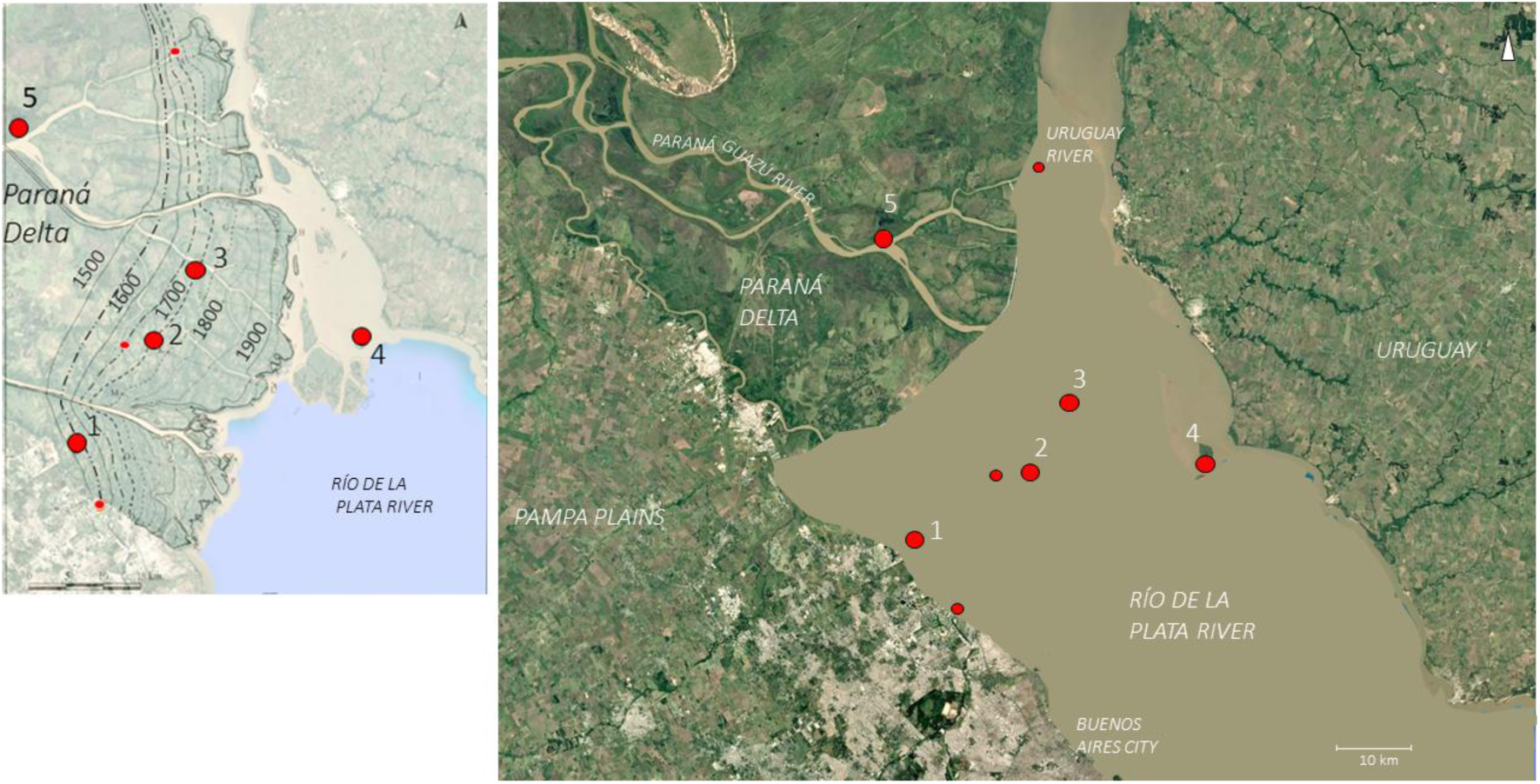
Left map: Location of the successive Paraná Delta advancing fronts according to Medina and Codignotto (2013) up to 1750 CE. The advancing fronts prior to 1750 CE are estimated in this study using isometric linear extrapolation, based on the most conservative progradation rates. Right map: Approximate location of the Paraná Delta’s advancing front around 1500 CE, along with the locations of currently known Guaraní sites. 1: Arroyo Malo – La Glorieta. 2: Arroyo Fredes. 3: Arroyo Largo. 4: Arenal Central. 5: Paraná Guazú 3 (The location of this site is approximate). Smaller red points represent isolated Guaraní burial urns as well as disturbed or destroyed Guaraní archaeological sites (see text).

Based on this overlapping evidence, it is possible to infer that Martín García Island was located at least 30–35 km from the Delta front in 1500 CE. However, it cannot be ruled out that several sedimentary islands had already formed between these settlements and the advancing front. In fact, the existence of numerous islands in the Río de la Plata estuary is supported by historical descriptions and maps, such as Levinus Hulsius’s map (ca. 1599), which suggests that the progradation process during the 16th and 17th centuries was characterized by the emergence of multiple small islands in the upper estuary. These islands were later consolidated into a more cohesive landmass, forming a more clearly defined Delta front, as depicted in later historical maps.

The insular nature of the known Guaraní sites in the area—each of them proably occupied prior to 1500 CE—closely aligns with the terminology used by early chronicles and Spanish colonial authorities to describe the Guaraní living near Buenos Aires, referred to as “Guaraníes de las islas” (“Guaraní of the islands”) (Fernández, 1582).

### 7.4. Catchment area and trade systems

Martín García Island has limited subsistence resources due to its small size, which would have required a population of forager-horticulturalists to consider a broader resource catchment area, including continental sectors of the estuary and the islands of the Paraná Delta. It is likely that crops were cultivated on the island, as it has suitable soils, evidenced by multiple plantations developed from historical times to the present. The aquatic areas surrounding the island also provided a notable abundance of fish. However, the possibility of cultivating plots in the riparian forest area of Martín Chico (Figure 21) cannot be ruled out. In fact, the procurement of certain faunal resources suggests that the riparian forest and adjacent steppe in that area were part of Arenal Central’s resource catchment zone (Section 6.3). Although open-environment species (e.g., *O. bezoarticus* and *R. americana*) identified in the site’s faunal assemblage could also have been hunted on the Argentine Pampas plains, this area is located 35 km in a straight line from the island, compared to just 3.5 km from the Uruguayan steppe.

**Figure 21.**
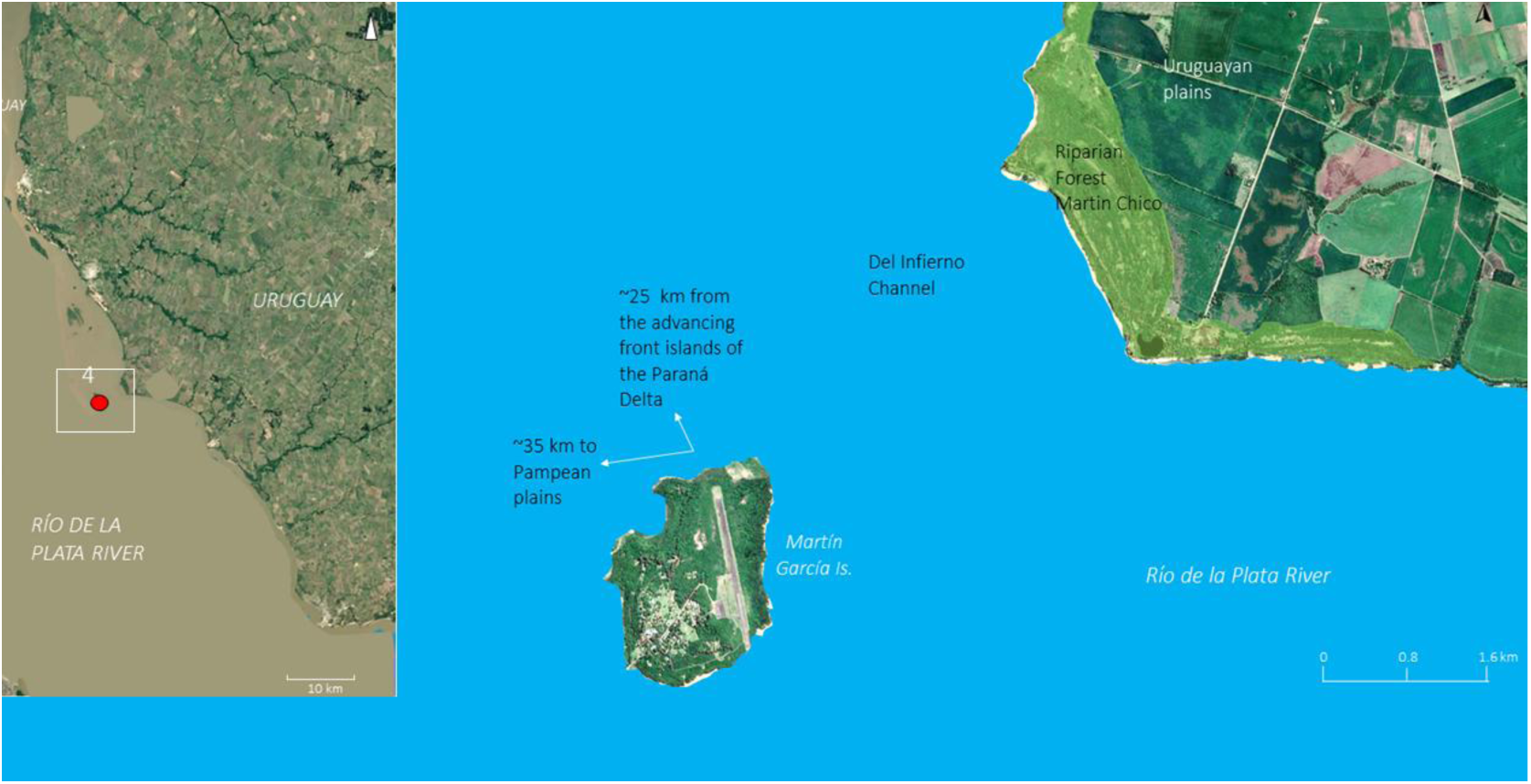
Left: location of Martín García island (as # 4). Right: Martín Chico area on the Uruguayan coast in relation to Martín García Island and distances to Paraná Delta and the Pampas plain at the time of the Guaraní occupation.

The lithic resources present at Arenal Central significantly expand the resource catchment area, this time toward the north of the island. The fluvial pebbles of cryptocrystalline and microcrystalline rocks recovered at the site are found on terraces and sandbars along the lower Uruguay River (Gentili et al., 1974; Gentili & Rimoldi, 1979; Silva Busso & Amato, 2017). In this area, basalts and silicified limestones are also available, and these have likewise been identified at Arenal Central. Silicified sandstones are also present in the area surrounding the lower Uruguay River, such as the piece shown in Figure 18-A, which likely corresponds to the Salto Chico Formation (Gentili & Rimoldi, 1979; Franco, 2014). Although these sandstones resemble those of the Ituzaingó Formation, which outcrop along the Paraná River and in inland areas of Entre Ríos (Silva Busso & Amato, 2017), the lithic assemblage from Arenal Central suggests that all of them originated from the lower Uruguay River. The acquisition of these rocks may have involved direct procurement, exchange, or a combination of both mechanisms, particularly considering the distribution of Guaraní sites in the estuary toward the lower Uruguay River.

Although chroniclers indicate that the Guaraní possessed metal artifacts acquired through their intra-ethnic network extending to the Andean region (Rodríguez, 1528 in Madero, 1939; Combès, 2008, 2015), metallic objects are extremely rare at Guaraní sites in the Delta. At Arroyo Fredes, two pure copper artifacts were recovered that could have an Andean origin, along with a third piece, an alloy likely of European origin (Buc et al., 2014). At Arenal Central, no metal artifacts have been recovered that can be definitively associated with the site’s occupation.

To conclude this section, it is worth highlighting that the bone point described in Section 6.5 (Figure 16-A) is not of Guaraní origin but instead exhibits traits typical of hunter-gatherers from the Paraná Delta and adjacent continental areas (Loponte, 2008; Buc, 2012). Its presence at Arenal Central is unlikely to result from exchange, as no artifacts associated with these populations have been found at the site. A plausible explanation is that it reflects episodes of interpersonal violence in the deltaic or nearby continental regions.

### 7.5. Guaraní occupations in the Paraná Delta and the Río de la Plata estuary

A recent reanalysis of the entire Guaraní radiocarbon dataset from southeastern South America suggests that the colonization of the Paraná Delta took place during Phase III of their expansion, which began around 1300 CE and extended to around 1600 CE (Loponte et al., 2025). This phase marks the peak of their geographical dispersal. In regions already occupied, a notable increase in the number of archaeological sites is observed beginning around 1300 CE—or even slightly earlier—suggesting demographic growth or the intensification of local settlement patterns. In contrast, in newly colonized areas such as the Paraná Delta, the Guaraní initially appear as a faint archaeological signal around 1300 CE, followed by a sharp increase in site frequency around 1490 CE (Figure 22). Even when applying a Bayesian averaging model to the radiocarbon dates from the Arenal Central and Arroyo Fredes sites—intended to mitigate potential oversampling biases— this marked upward trend remains clearly visible (Figure 22). These findings suggest an initial pre-Columbian colonization by a small founding population with low demographic density, followed by a significant population increase shortly before the onset of the historical period. This demographic surge may reflect the arrival of new groups in the region, internal population growth, or a combination of both.

**Figure 22.**
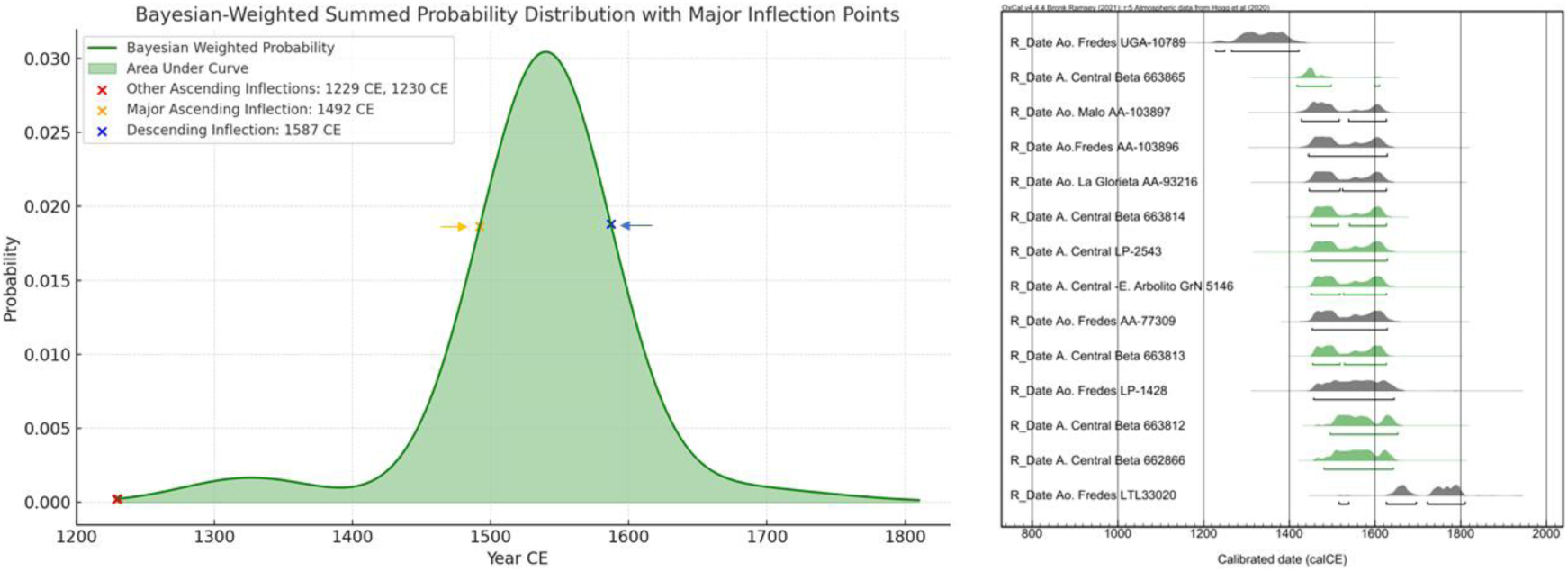
LEFT graph: Bayesian-weighted summed probability curve for radiocarbon dates from Guaraní sites in the Paraná Delta and Río de la Plata estuary. The curve was smoothed using a Gaussian filter (sigma = 10), and inflection points were identified through the second derivative. Right: calibrated ranges calculated using OxCal 4.4.4 (Bronk Ramsey, 2021) and SHCal-20 (Hogg et al., 2020). The green ranges correspond to Arenal Central – El Arbolito, as referenced in Table 1 of this work. The dates for Arroyo Fredes were taken from Loponte et al. (2011a), except for AA-103896 (Bonomo et al. 2015), as well as the radiocarbon ages from Arroyo La Glorieta and Arroyo Malo sites. The Arroyo Fredes LTL 33020 radiocarbon determination was taken from Loponte et al. (2024).

This pattern aligns with models of pulsed migration and frontier expansion (e.g., Lightfoot and Martinez, 1995), where small pioneering groups establish an initial presence, often with low archaeological visibility, later followed by more substantial population movements that consolidate the occupation. Such a process may reflect strategic mobility, adaptive responses to ecological opportunities, or social mechanisms of territorial incorporation and alliance-building with new arriving populations.

Just as Guaraní colonization appears as a sudden event within an archaeological timescale, it also declines abruptly starting in 1587 CE (Figure 22), approximately contemporaneous with the second foundation of Buenos Aires in 1580 CE. This city quickly established itself as a colonial administrative center and an overseas port. Furthermore, in 1582, the *encomienda* system was implemented, integrating indigenous populations around Buenos Aires.

As part of this allocation of indigenous people to *encomenderos^4^*, 14 Guaraní *caciques* (chiefs) were included along with the population under their command (Fernández, 1582), suggesting that at least 14 Guaraní villages existed around Buenos Aires at that time. The proximity of these villages to the newly re-founded city, their predominantly insular situation, and the probable Spanish interference or destruction of the intra-ethnic networks of a population like the Guaraní— with stable villages dependent on large exploitation territories and their social connections— likely had a highly negative impact on their way of life, making them among the first Indigenous communities in the area around Buenos Aires to become disarticulated.

The Guaraní colonization of the Paraná Delta and Río de la Plata estuary region is unique since they predominantly used the islands to establish their villages, forming what appears to be a roughly linear pattern parallel to the Delta’s advancing front (Figure 20). This differs significantly from settlement patterns in the northern continental sectors, showcasing the Guaraní’s adaptability to a new environment characterized by insularity and its environmental constraints. However, it seems that the choice of this suboptimal insular settlements was a consequence of resistance from local groups to Guaraní penetration into the Delta and the whole Paraná wetland. These pre-existing non-Guaraní populations were composed of complex hunter-gatherers with some degree of ancillary horticulture, high demographic density, and, above all, well-developed active defense strategies for productive territories that emerged around 1100 CE and likely intensified following the arrival of the Guaraní (Loponte et al., 2006; Acosta & Loponte, 2013). This resistance may have taken the form of various local alliances to counter Guaraní expansion. Indeed, early European chronicles document “permanent” conflicts between local societies and the Guaraní, distinguishing these two major population groups as being in constant competition (e.g., Ramírez, 1528, in Madero, 1939; Fernández de Oviedo y Valdés [ca. 1526–1557] 1945).

Given these social constraints, occupying the Río de la Plata estuary—such as in the case of Arenal Central—may have represented a viable alternative for Guaraní expansion. However, Guaraní colonization of the estuary remained relatively limited. To date, the only identified site potentially representing a Guaraní settlement beyond Martín García is located at Arroyo Las Cañas, 70 km south of Arenal Central, on the right bank of the intermediate Río de la Plata estuary near the city of La Plata (Maldonado Bruzzone, 1931) (Figure 23). This area also featured a discrete sector with riparian forest (such as Martín Chico, across from Martín García), a small remnant of which still exists today. Additionally, the encomienda list from 1582 CE suggest a faint presence of Guaraní groups in the middle portion of the estuary (Fernández, 1582).

**Figure 23.**
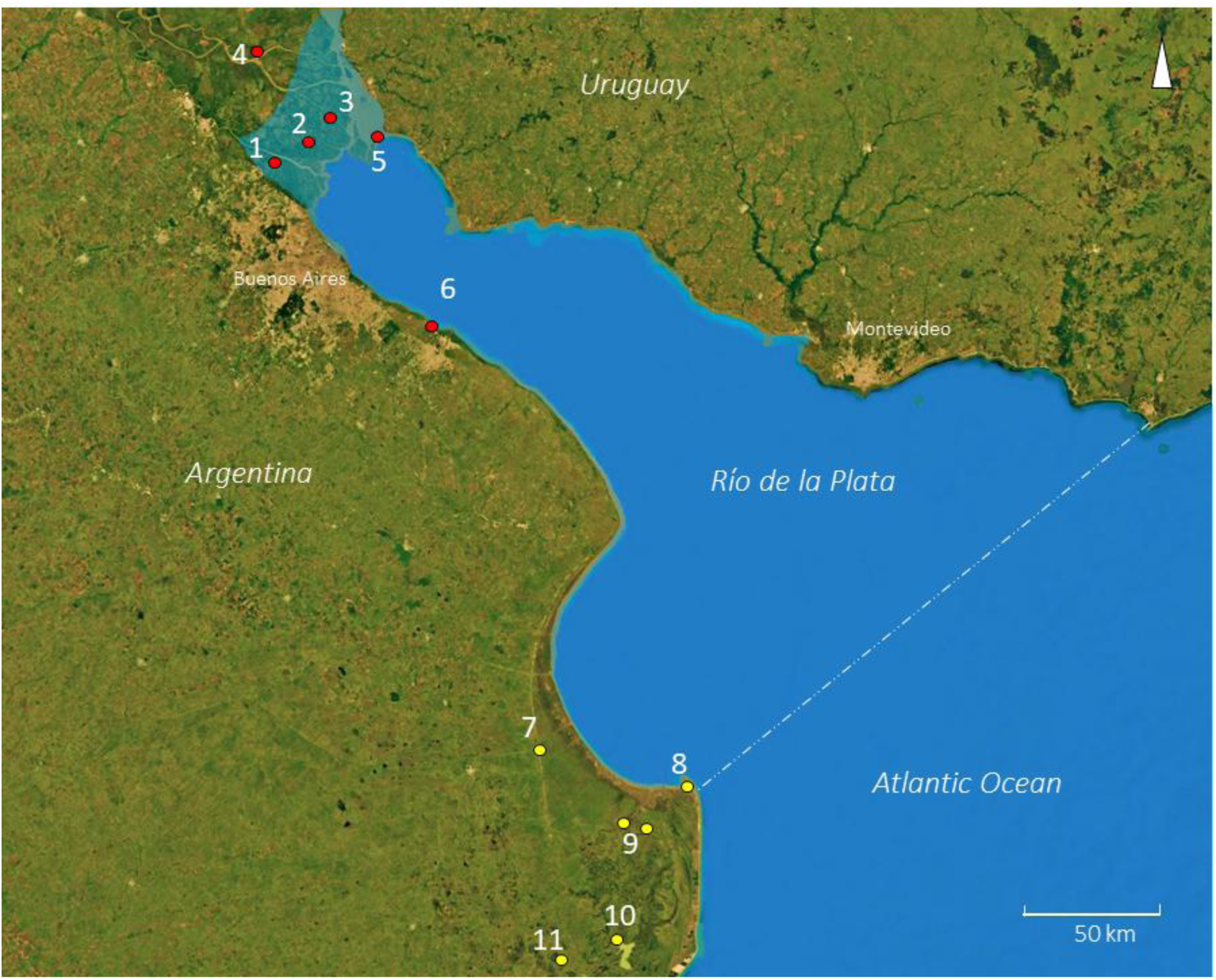
Locations of Guaraní sites in the Paraná Delta and the Río de la Plata estuary based on the most likely position of the Delta front around 1500 CE: 1 = Arroyo Malo & La Glorieta. 2 = Arroyo Fredes. 3 = Arroyo Largo. 4 = Paraná Guazú 3 (there is no precise information about the location of this site). 5 = Arenal Central (Martín García island). 6 = Las Cañas (uncertain assignment, see text). In yellow, examples of hunter-gatherer sites with Guaraní pottery: 7 = Los Molles. 8 = Divisadero 6. 9 = Canal 2 & La Loma. 10 = La Isolina. 11 = La Zeta (Aldazabal & Eugenio, 2013b).

This limited expansion along the estuary may have been influenced by two factors. The first relates to less favorable environmental conditions for Guaraní niche construction. Unlike the lower Paraná Delta and the entire Paraná wetland, which is characterized by extensive riparian and marginal forests covering hundreds of square kilometers, the estuary features small, isolated patches of subtropical forest along the riverbanks, with open grasslands beginning almost immediately beyond the shores. The second factor could be risk minimization, given the successive arrival of Spanish and Portuguese expeditions starting in 1516 CE and the establishment of settlements along the lower Paraná and Uruguay rivers, as well as in the estuary itself, which may have discouraged or halted the expansion process.

Along the estuary, extending to the lower course of the Salado River and even beyond to the area where the Atlantic Ocean begins, small quantities of Guaraní ceramics have been found within non-Guaraní contexts (e.g., Aldazabal et al., 1995; Aldazabal & Eugenio, 2013a, 2013b; González & Frère, 2013) (Figure 23). These findings consist of only a few fragments within sites generated by local hunter-gatherers. Guaraní ceramics are also scattered across a large number of sites in the Paraná and Uruguay rivers. However, these are not Guaraní sites *per se*, but rather locations where their pottery is present—possibly as a result of exchange.

Guaraní pottery may have been prestige items among local populations, entering exchange networks as prized trade goods. Thus, Guaraní vessels likely arrived through the circulation of goods across the estuary via long-distance exchange or individual mobility, rather than representing permanent Guaraní settlements like those in the Paraná Delta or Arenal Central. Extensive research by González, Frere and colleagues in the Salado Depression, and by Aldazabal and Eugenio along the southern Atlantic coast, rules out sampling bias as the reason for the absence of Guaraní sites. Although a Guaraní village may yet be discovered, none has been identified to date.

## 8. Final Remarks

Arenal Central was occupied at a later stage following the initial Guaraní colonization of the region, likely representing the southernmost settlement of the Amazonian forager-horticulturist expansion along the Uruguay River axis. As such, it may reflect an attempt to expand further into the rest of the Rio de la Plata estuary, given the social constraints on the colonization of the Paraná Delta.

Arenal Central functioned as a residential base for several decades, with its initial occupation beginning in the pre-Columbian period and possibly being abandoned at the onset of the historical times. The site’s archaeological record exhibits typical characteristics of Guaraní residential sites, including vessel types, decoration, surface treatment, paste composition for ceramic manufacturing, lithic and bone artifact types (with some previously noted exceptions), as well as faunal exploitation strategies similar to those observed at other Guaraní sites.

Some variations in the archaeological record are also noted compared to other sites, such as a slight impoverishment or simplification in ceramic decorative styles, although this may be a sampling bias that requires further collection expansion. Due to the settlement’s insular nature and the small size of the island, the population utilized a broad resource acquisition area based on fluvial mobility, including the exploitation of open grasslands, demonstrating a significant degree of adaptability to new environments. Ongoing research will expand the available information on this site and its role within the broader context of Guaraní colonization in the region.

## 9. Acknowledgments

Special thanks to Nazareno Asin, Gloria Domínguez, and all the members of the ranger team at the Martín García Natural Reserve for their essential support during our fieldwork on Martín García Island.

## Footnotes

1 This work includes 119 bibliographic references, ⁓ 90 of which correspond to studies conducted by ⁓ 140 researchers other than the authors of the present study. We trust that readers will be understanding regarding the number of cited works, as each was selected for its direct relevance or the quality of information it provides in relation to the subject of this research.

2 Collections obtained unsystematically, small collections, and those from highly impacted sites were not considered due to the different biases they present.

3 Several historical records provide fairly consistent information suggesting that at least two earlier expeditions took place; however, their existence is not unanimously accepted among scholars (e.g., Levillier, 1948; Madero, 1939), a topic we will address elsewhere.

4 The *encomenderos* were settlers granted the right to collect tribute and labor from indigenous people, whom they were also required to protect.

